# Mitochondrial DNA copy number in neurodegenerative diseases: a global meta-analysis of 156 comparisons across 76 studies

**DOI:** 10.64898/2026.08.25.747144

**Authors:** Reuben Mathews, Sifallah B. Bouyadjera, Jennifer J. Donegan, Justin C. Havird

## Abstract

Mitochondria are central hubs for cellular metabolism and mitochondrial dysfunction is a hallmark of many chronic diseases. Consequently, changes in mitochondrial DNA copy number (mtDNA-CN), the number of mtDNA genomes per cell or tissue sample, are associated with diseases ranging from cancer and obesity to psoriasis and all-cause mortality. MtDNA-CN especially holds promise as a biomarker for neurodegenerative diseases given the high-energy demands of neurons, but the relationship between neurodegeneration and mtDNA-CN is controversial due to mixed results among studies. Here, we performed a systematic review and meta-analysis to examine how mtDNA levels are altered in neurodegenerative diseases with the goal of identifying potential moderators that explain variation among studies. After systematic review, 76 studies met our inclusion criteria, yielding 156 comparisons between mtDNA-CN in control and disease populations. Overall, mtDNA-CN was ∼5.3% lower with neurodegeneration, but the difference was not statistically significant (14% decrease – 4% increase, *P* = 0.244), with extreme heterogeneity among studies (*I*^2^ = 99.5%). Results varied significantly among neurodegenerative diseases, with Alzheimer’s showing a convincing 21% decrease in mtDNA-CN, but no change in mtDNA-CN in Parkinson’s despite large sample sizes. Decreases in mtDNA-CN with neurodegeneration were also more extreme at older ages. Surprisingly, the tissue and assay method used to quantify mtDNA-CN did not influence overall patterns. However, significant interactions were found among moderators. For example, mtDNA-CN decreased in cerebrospinal fluid in Parkinson’s, but not for Alzheimer’s. Studies that were published in earlier years also showed more extreme decreases in mtDNA-CN with neurodegeneration, while more recent studies tended to have modest effects. While our analyses explained some variation among studies, excessive heterogeneity persisted even after accounting for all moderators and their interactions (*I*^2^ = 85.7%). We conclude that the general perception of decreased mtDNA-CN with neurodegeneration is a vast oversimplification that may stem from “legacy” effects of early studies. However, mtDNA levels offer great promise as biomarkers for neurodegeneration, other diseases, and general health metrics, assuming appropriate complications are considered.

**Significance statement:** Mitochondrial dysfunction is associated with many diseases and the number of mitochondrial genome copies in a given (mtDNA-CN) is a promising biomarker for overall health and disease risk. While mtDNA-CN has been linked to neurodegenerative diseases, many studies find contradictory results. Here, we performed a systematic review and meta-analysis of 76 studies compromising 156 comparisons of mtDNA-CN in populations with and without a neurodegenerative disease. Overall, we found only a small, statistically non-significant decrease in mtDNA-CN with neurodegeneration, but results varied widely among studies. The diagnosed disease (e.g., Alzheimer’s vs. Parkinson’s), age of individuals, and date of the study explained some of this variation, while methodological differences did not. We provide a systematic appraisal of when mtDNA-CN may be useful as a biomarker for neurodegeneration and future studies needed for further evaluation.

## Introduction

Mitochondria are central hubs of cellular metabolism. They are primarily known for energy production through oxidative phosphorylation (OXPHOS), but also play roles in fatty acid/cholesterol synthesis, generation of cellular reactive oxygen species (ROS), regulation of cytosolic calcium, and programmed apoptosis, among others. Consequently, disruptions in mitochondrial function are associated with many diseases (1–3), particularly those that affect the central nervous system (4, 5).

The brain consumes a disproportionate amount of energy and neurons are extremely “energy hungry” cells that require a continuous flow of ATP to sustain physiological processes (e.g. action potential firing, synaptic transmission, and maintenance of ionic gradients) that underlie complex functions like sensation, motor control, and cognition (6, 7). Because mature neurons get around 93% of their ATP from OXPHOS, they are vulnerable to disturbances in mitochondrial function (8). Not surprisingly, mitochondrial dysfunction contributes to the pathophysiology of many neurodegenerative diseases, including Parkinson’s disease, Alzheimer’s disease, amyotrophic lateral sclerosis (ALS), multiple sclerosis (MS), and others (9). For example, in Parkinson’s disease mitochondrial function is impaired in idiopathic forms of the disease, loss-of-function mutations in the mitochondrial quality control gene parkin (*PRKN*) are associated with autosomal recessive forms of the disease, and drugs that inhibit OXPHOS increase disease risk (10). In Parkinson’s, changes in mitochondrial function can lead to impaired mitophagy, reduced ATP production, oxidative stress, and ultimately the degradation of dopaminergic neurons in the substantia nigra pars compacta. While an important contributor to neurodegeneration, assessing mitochondrial health and developing mitochondrial biomarkers for neurodegenerative diseases is not straightforward.

Mitochondrial DNA copy number (mtDNA-CN) is an emerging biomarker for mitochondrial health, quantifying the number of mtDNA genomes per cell or tissue sample (11). MtDNA-CN is predictive of many diseases (11–14) and reduced mtDNA-CN can be associated with lower abundance of mitochondria and compromised mitochondrial maintenance, ultimately leading to reduced mitochondrial/OXPHOS function (15, 16). Paradoxically, elevated mtDNA-CN can also be interpreted as a compensatory mechanism for mitochondrial dysfunction or impaired OXPHOS, as a nuclear response to dysfunctional mitochondria may be to increase the number of OXPHOS proteins via increased replication of mtDNA (17, 18). However, mitochondrial function can also change independent of mtDNA-CN via changes in mtDNA transcription, translation, post-translational modifications, or other regulatory processes. For example, Parkinson’s disease lymphoblasts were reported to have normal mtDNA-CN levels, despite significantly elevated mitochondrial respiration and ATP synthesis (19).

Not surprisingly, mtDNA-CN has been assessed across a range of neurodegenerative diseases (20–23). While the general consensus may be that “more is better” when it comes to mtDNA-CN and human health (24), findings from neurodegenerative diseases have been highly inconsistent. For example, in Parkinson’s disease, different studies report reduced, increased, and unchanged mtDNA-CN (12, 25–27). Similar contradictory results have been reported for Alzheimer’s disease, ALS, and MS (28–32). Overall, these findings suggest that mtDNA-CN may not behave as a uniform biomarker for neurodegenerative disorders, but may show variable patterns moderated by many factors, including the tissue sampled, type of disease, age of patients, and the method of quantification (33, 34). Although prior reviews have highlighted these discrepancies among studies, they have been confined to specific tissues or diseases and conclude that associations between mtDNA-CN and neurodegeneration are “controversial” (21, 35). Formal meta-analyses of mtDNA-CN during neurodegeneration are limited to a single study to our knowledge, based on only 13 studies and focused solely on Parkinson’s disease (35). Overall, these findings suggest mtDNA-CN could be a useful biomarker for neurodegenerative diseases, but systematic sources of variation among individuals must first be investigated.

Here, we conducted a comprehensive systematic review and meta-analysis of published studies that quantified mtDNA-CN in individuals with neurodegenerative diseases compared to healthy controls. We aimed to identify overall trends and characterize the perceived heterogeneity in the literature. We hypothesized that mtDNA-CN would be reduced in individuals with neurodegenerative disease relative to controls, but also that the magnitude and direction of this association would differ substantially across studies due to biological and methodological moderators. Specifically, we evaluated whether tissue source, disease diagnosis, underlying neuropathology, age, geographic location, and method of mtDNA-CN quantification affected how mtDNA-CN changed with disease. By combining dozens of primary studies, we hoped to identify trends that would be impossible to investigate in individual studies, ultimately informing how and when mtDNA-CN could serve as a useful biomarker for neurodegeneration.

## Materials and methods

### Systematic literature search

A systematic literature search (Table S1) was conducted in early 2025 to identify studies that quantified mtDNA-CN in individuals with diagnosed neurodegenerative diseases compared to non-diseased controls. Searches were conducted using three databases: PubMed, Embase, and Web of Science. Additionally, the first 100 entries from a Google Scholar search were examined to ensure a broad breadth of coverage across the biomedical literature.

The primary strategy for developing search strings combined terms related to mitochondria, mtDNA copy number, and neurodegenerative disease types (Table S1). Core search terms included combinations of mitochondri* AND (“copy number” OR “CN” OR “mtDNA-CN” OR “Mitochondrial DNA Copy Number”) AND disease-specific terms such as Parkins* OR Alzheim* (Table S1). Database specific syntax (e.g., MeSH terms) was used where appropriate.

Studies returned from these searches were imported into Rayyan (36) for deduplication and screening. During screening, studies were retained if they met the following inclusion criteria: were primary research articles published in peer-reviewed journals in English, examined human populations with a diagnosed neurodegenerative disease and an appropriate non-disease control group, quantified mtDNA-CN, and reported mtDNA-CN for both groups in a way where appropriate data (means, sample sizes, and variances for both groups) could be extracted or extrapolated. Most studies normalized mtDNA content to at least one single-copy nuclear reference gene, allowing for estimation of the normalized number of mtDNA genomes per cell (i.e., the mitochondrial-to-nuclear DNA ratio). However, some studies did not use a nuclear gene for normalization, especially those that quantified cell-free mtDNA (e.g., as mtDNA copies per microliter of plasma). Studies were excluded if they were conference abstracts without an associated peer-reviewed manuscript, were systematic reviews, were case studies of single individuals, did not include a control population, or did not report mtDNA-CN in a way where relevant data could be extracted. We also excluded one study where a sample size of one was used for the disease population (37).

The screening process occurred in two stages. First, after deduplication two authors (RM and JCH) screened studies based primarily on their titles and abstracts to remove clearly irrelevant studies (e.g., reviews, non-primary studies, or those not involving neurodegeneration). To evaluate screening consistency, 15% of the studies were randomly selected and independently re-screened by the other author. Reviewer agreement between RM and JCH was high, as quantified using Cohen’s kappa statistic (raw agreement = 94%, kappa = 0.63) (38). After initial screening, all authors screened remaining studies based on the full text following the pre-defined inclusion and exclusion criteria. Studies were also included based on forward and backward literature searches of articles that met our inclusion criteria. During the screening process, two relevant systematic reviews/meta-analyses were published (21, 35), which were also used as sources for relevant primary studies.

### Data extraction

For all included studies, mtDNA-CN data were extracted from disease and control groups, including means, measures of variance (converted to standard deviations when other metrics were provided), and sample sizes. When numerical values were not reported directly in tables or the text itself, data were extracted from figures using WebPlotDigitizer 4.2. We generally followed how effect sizes were reported in the original articles when extracting data. For example, if authors originally reported separate effect sizes for males and females (39) or for idiopathic vs. familial ALS (40), we recorded separate effect sizes. One exception was for Lunnon et al. (2017) (41), which reported 12 separate effect sizes for 12 different mtDNA markers. In that case, we only extracted effect sizes for two different mt markers (*ND1* and *CYTB*, the two most popular markers based on other studies).

Metadata were also recorded for each study, including both organizational details and potential moderators to explain variation among studies. These included the article and effect size number, year of publication, the author who extracted the data, the data extraction date, the data’s source in the article, and the original measure of variance. Variables extracted as potential moderators included tissue type used to calculate mtDNA-CN, the diagnosed neurodegenerative disease, average age of control and disease populations (and their difference), whether control and disease populations were age-matched (either as stated by the original authors or based on reported average ages being < 10 years different), sex distribution (relatively matched – either as reported by authors or based on reported sex ratios being < 10% different, and male-or female-biased), geographical population (recorded as country), assay method (i.e., qPCR, dPCR, or sequencing based), the number of mt and nuclear markers used to calculate mtDNA-CN, and which regions/genes of mtDNA and nuclear DNA were used.

Based on how authors reported moderators, we consolidated some groups to facilitate meta-analyses. For tissue type, consolidation yielded the following categories: whole blood, cellular blood fraction (mainly leukocytes), acellular blood fraction (mostly plasma or serum), cerebrospinal fluid, brain tissue, peripheral tissue (fibroblasts or muscle), and cell lines (all were lymphoblastoid cell lines). For disease, two moderating categories were established: the first was based on diagnosed disease and included amyotrophic lateral sclerosis (ALS), Alzheimer’s disease, dementia, Huntington’s disease, mild cognitive impairment (MCI), multiple sclerosis (MS), Parkinson’s disease, and an “other” category that included several diseases with limited numbers of effect sizes (e.g., Machado-Joseph disease and Aicardi-Goutières syndrome). Diseases were also categorized using a second moderator category to account for underlying neuropathology and included the following categories: demyelinating disorders, polyglutamine/CAG repeat diseases, proteinopathies, synucleinopathies, tauopathies, and an “other” category (e.g., prion diseases and all-cause dementia).

Geographical location was largely consolidated to reflect continental differences, although the United States and United Kingdom had many data points, so were treated as separate locations (Australia and Africa were only represented by single data points and were not included in this moderator analysis).

### Meta-analyses

Meta-analyses were conducted in R 4.3 using the metafor package (42). Effect sizes were calculated for each pair of mDNA-CN values comparing a disease vs. control population as the natural logarithm of the response ratio (ln *RR*) using the escalc function in metafor. For this effect size metric, values less than zero indicate lower mtDNA-CN with neurodegeneration, while values greater than zero indicate higher mtDNA-CN with neurodegeneration. In some instances, we report the untransformed response ratio to increase clarity (e.g., the % increase or decrease in mtDNA-CN with neurodegeneration), but all statistical analyses were performed on ln *RR* values.

Meta-analyses were conducted using mixed-effects/random models using rma and rma.mv functions in metafor. Because multiple effect sizes were often calculated within the same article (e.g., for different tissues, different markers, or using different populations), publication was included as a random effect in most models. Heterogeneity in meta-analytical models was assessed using *I*^2^ and *Q* statistics (with associated *P* values for the latter). After conducting analyses with no moderators to generate overall effect sizes, we then performed meta-analyses with single moderators (e.g., tissue or disease, not both) to examine overall trends and importance of moderators in explaining heterogeneity in effect sizes. For the moderators that explained a significant amount of heterogeneity, their importance was ranked based on AIC values using a smaller dataset where information was available for all moderators of interest (e.g., studies where age could not be extracted were excluded). Most moderators were categorical, but age was treated as a continuous moderator (i.e., in a meta-regression model) based on the average age of the disease population. For moderators that explained a significant amount of heterogeneity, we investigated models that included interactions between two or more moderators. For example, we asked whether differences in mtDNA-CN were more extreme among tissues in Parkinson’s vs. Alzheimer’s disease. We also investigated whether moderators were confounded with each other using chi-squared tests for categorical moderators and investigating interaction plots using continuous moderators. Finally, we used a model including all significant moderators and their interactions to determine how much heterogeneity could be explained after accounting for all identified sources.

Publication bias was assessed by manually examining a funnel plot for asymmetry and performing Egger’s regression (43), trim-and-fill analysis (44), and a fail-safe number analysis (45). We also performed a cumulative meta-analysis (46) to assess how stable effect sizes were over time. Finally, we performed sensitivity analyses, including “leave 1 out” analysis to determine stability of the main results. Figures were created using the orchard and ggplot2 packages in R (47, 48).

### Data availability

All data for our analyses are publicly available via FigShare (link will be provided after acceptance – all relevant files have been made available to reviewers). This includes extracted data and metadata for the articles included in our analyses, calculated effect sizes, and R code used to produce the analyses and statistics reported here (most data are also available via supplementary Tables S1-S7).

## Results

### Popular characteristics of studies that quantify mtDNA-CN during neurodegeneration

Our literature search returned 669 unique articles for screening (Fig. S1). Ultimately, 76 studies met our inclusion criteria (see Methods) and were used for data extraction and meta-analysis, with publication years from 2000 to 2025 (Fig. S1, Table S2). We calculated a single effect size from most studies, comparing mtDNA-CN between disease vs. control populations as the natural logarithm of the response ratio (ln *RR*), although some studies produced many effect sizes due to examining multiple populations, diseases, tissues, or mtDNA markers (28, 34, 49). In total, 156 effect sizes were calculated. Parkinson’s and Alzheimer’s diseases were represented most in our dataset (*k* = 64 and 33 effect sizes, respectively). Blood-based tissues, brain tissue, and cerebrospinal fluid were the most popular tissues examined, ranging from *k* = 18 effect sizes in the acellular fraction of blood to *k* = 46 effect sizes in the cellular fraction. Studies from populations in East Asia, Europe, the United Kingdom, and the United States were most common, making up ∼94% of the calculated effect sizes. Of the studies that reported sex ratios for their populations, the vast majority (77%) had relatively matched sex ratios with similar numbers of sexes in both disease and control populations and did not report sex-specific data. Similarly, for 138 out of 154 effect sizes where age was reported, ages of disease and control populations were relatively matched (either as stated by authors or < 10 years difference). Across all studies, average ages of control and disease populations were 64.4 years (range: 6 – 89 years). Most studies (79%) used a single mtDNA marker and regions of *ND1*, the D-loop, and *CYTB* were popular, although for 20 effect sizes sequencing methods were used and reads were mapped to the entire mtDNA to calculate mtDNA-CN. mtDNA-CN was normalized to at least one nuclear marker in 126/156 studies, although studies that did not normalize mtDNA-CN to nuclear DNA were found across most diseases and tissue types. Finally, qPCR (*k* = 107), dPCR (*k* = 28), and sequencing-based approaches (*k* = 20) were used exclusively to calculate mtDNA-CN, although the earliest study in our dataset used southern blots (50).

### Overall, modestly lower mtDNA-CN during neurodegeneration with extreme heterogeneity among studies

We ran several meta-analytical models on the 156 effect sizes without including moderators to investigate the overall effect of neurodegeneration on mtDNA-CN (Table S3). The most appropriate model was determined *a priori* to include random effects among effect sizes, weighting effect sizes by precision (i.e., sample sizes within a study), and including a random effect of publication to account for studies that reported multiple effect sizes. In this model, overall mtDNA-CN was ∼5.3% lower in neurodegeneration populations compared to control populations (Fig. 1, ln *RR* = −0.054), but the difference was not statistically significant (range ln *RR* = −0.146 – 0.037, *P* = 0.244). However, heterogeneity was extreme among studies (from a 75% decrease to a 228% increase in mtDNA-CN with neurodegeneration, *I*^2^ = 99.5%, *Q* = 4680.7, *P* < 0.001). In other meta-analytical models, overall effects remained modest, although sometimes statistically significant (e.g., when using fixed effects), but heterogeneity was always high (*I*^2^ > 97% in all analyses, Table S3).

**Fig. 1.**
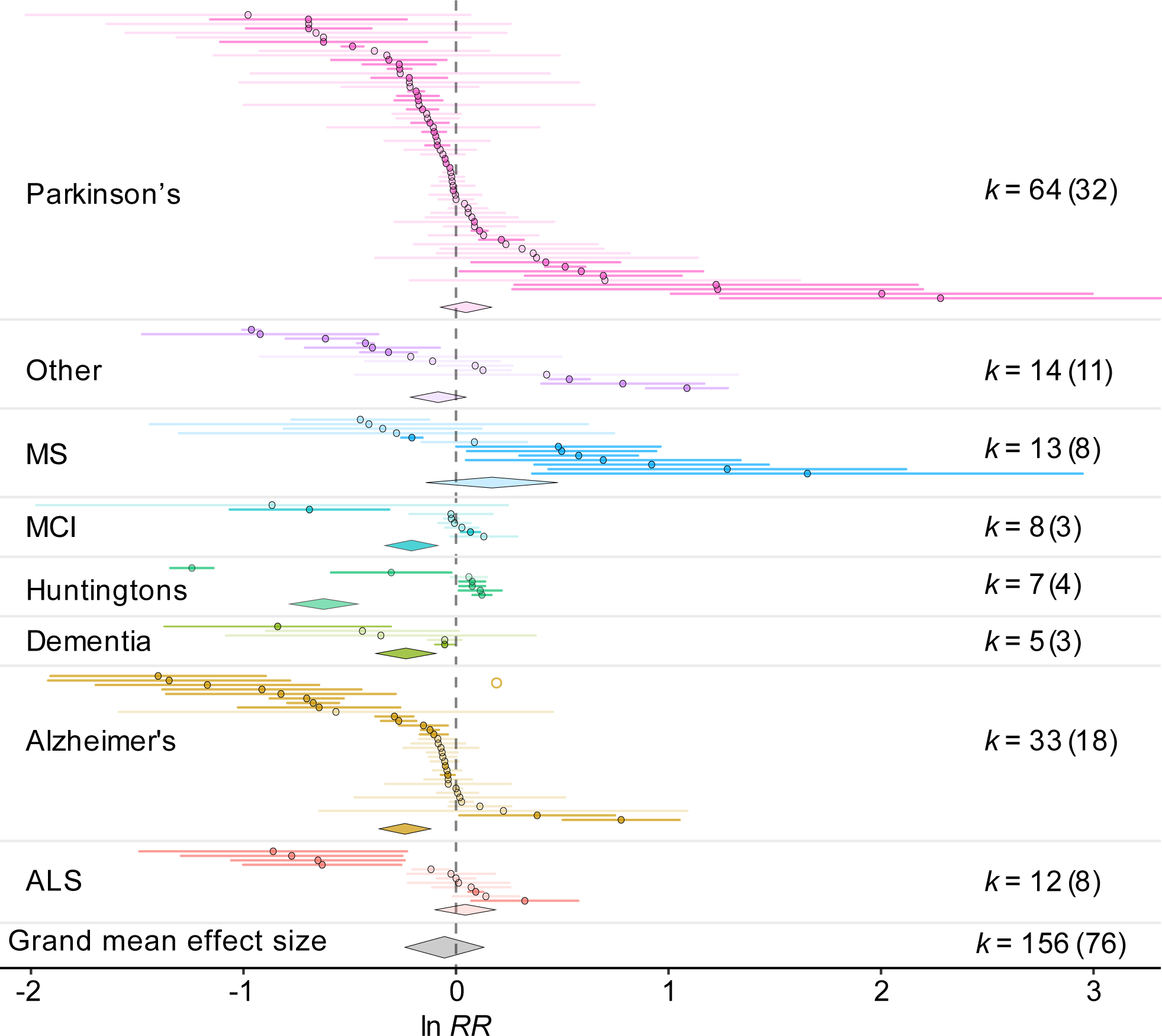
How mtDNA-CN changes with neurodegeneration varies among different neurodegenerative diseases. Effect size is displayed as ln *RR* on the x-axis, with values < 0 indicating lower mtDNA-CN with neurodegeneration and values > 0 indicating higher mtDNA-CN with neurodegeneration compared to control populations. Effect sizes are grouped and colored based on disease (as diagnosed) and ordered from lowest to highest for each disease. Mean and 95% confidence intervals are displayed per effect size. Darker colors indicate effect sizes with 95% CIs that do not overlap 0 while lighter colors do overlap 0. Overall grand mean effect sizes are colored similarly and displayed as diamonds for each disease, centered on the mean effect size with vertices indicating 95% CI. The grand mean effect size is displayed below all others. *k* values show number of effect sizes and the corresponding number of studies (publications) for each disease category.

### Disease, age, and sex (but not tissue) explain significant heterogeneity in effect sizes

Several of the moderators we investigated explained a significant amount of heterogeneity in effect sizes. We first explored individual moderators separately in models that included publication as a random effect (Table S4). Disease explained a significant amount of heterogeneity, both based on diagnosis (Fig. 1, *P* < 0.001) and when grouped according to neuropathology underlying the disease (Fig. S2, *P* < 0.001). Alzheimer’s patients showed a significant decrease in mtDNA-CN compared to controls (21% ± 6% decrease, *k* = 33 effect sizes, *n* = 18 studies), and similar results based on smaller sample sizes were found for dementia, MCI, and Huntington’s disease. Other diseases had no statistical difference in mtDNA-CN compared to controls, including Parkinson’s disease, which had the largest sample size in our study (5% ± 6% increase in mtDNA-CN with disease, *k* = 64 effect sizes, *n* = 32 studies).

Sex also explained significant heterogeneity in the dataset (Fig. 2A, *P <* 0.001). Studies with relatively matched proportions of males and females showed a modest, statistically nonsignificant decrease in mtDNA-CN with disease, similar to the overall effect size. However, studies that measured female- or male-biased populations showed a significant increase in mtDNA-CN with disease (25% ± 7% for females and 23% ± 6% for males).

**Fig. 2.**
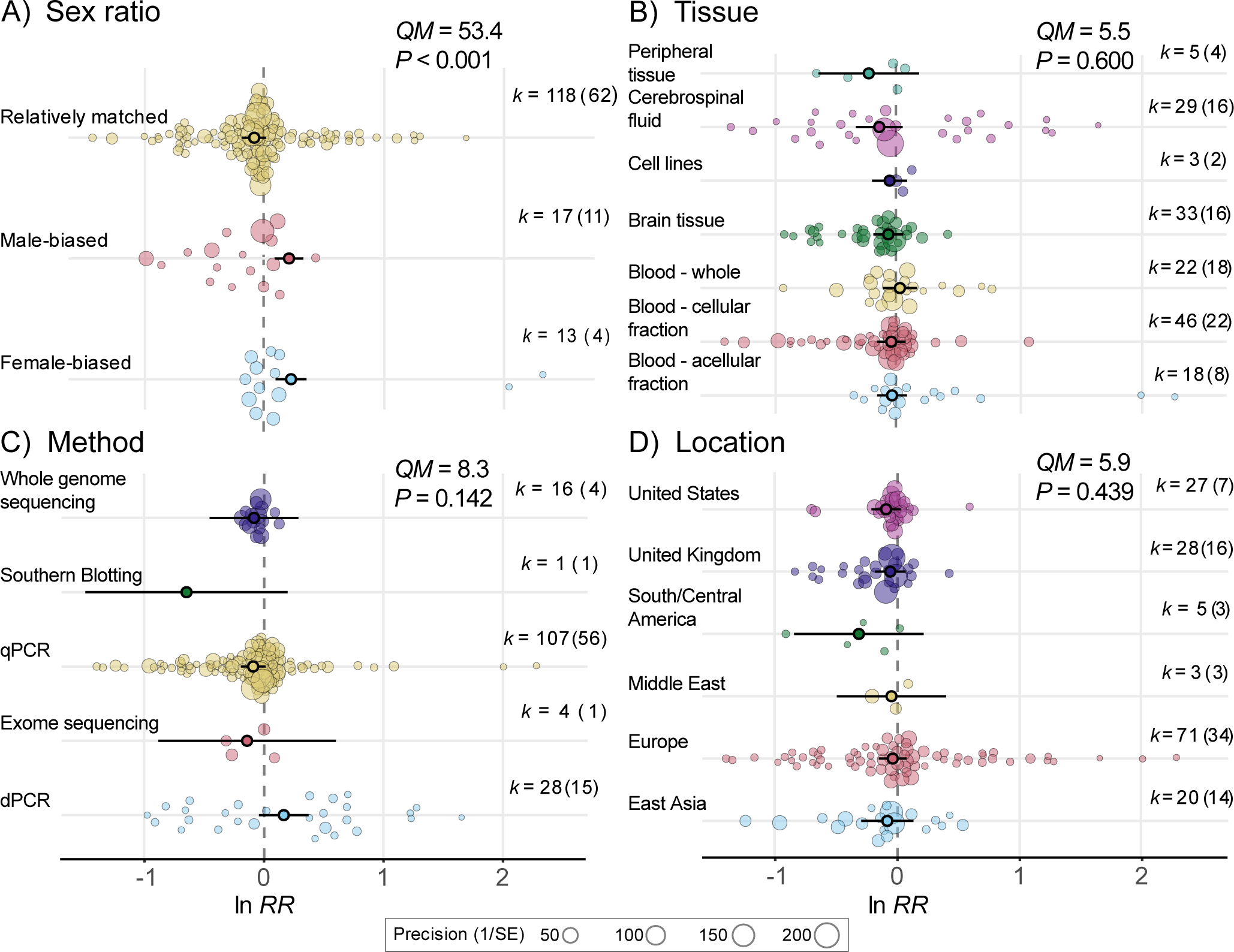
Moderators that do or do not explain why mtDNA-CN changes differently with neurodegeneration. A) The sex ratio of the populations explained significant heterogeneity among effect sizes, but the B) tissue examined, C) method of quantifying mtDNA-CN, and D) location of the study did not. Effect sizes are displayed on the x-axes as in Fig. 1, along with *k* values. Transparent points represent individual effect sizes and are scaled to their precision (i.e., sample sizes), while opaque points represent mean effect sizes per category and are shown with 95% CIs. *QM* and *P* statistics are displayed from mixed-effects meta-analytical models examining whether single moderators explain significant heterogeneity in effect sizes.

Population age also explained significant heterogeneity in the dataset (Fig. 3, *P* < 0.001). As population age increased, the mtDNA depletion in the disease population became more pronounced, equating to ∼5% lower mtDNA-CN in the disease population relative to the control population for every 10-year increase in age. For example, at age 50 mtDNA-CN would be about 2% higher in the disease population, but at age 80 mtDNA-CN would be about 14% lower in the disease population based on our dataset. A similar effect of age was found when considering either the age of the disease or control population (Fig. 3, Fig. S3).

**Fig. 3.**
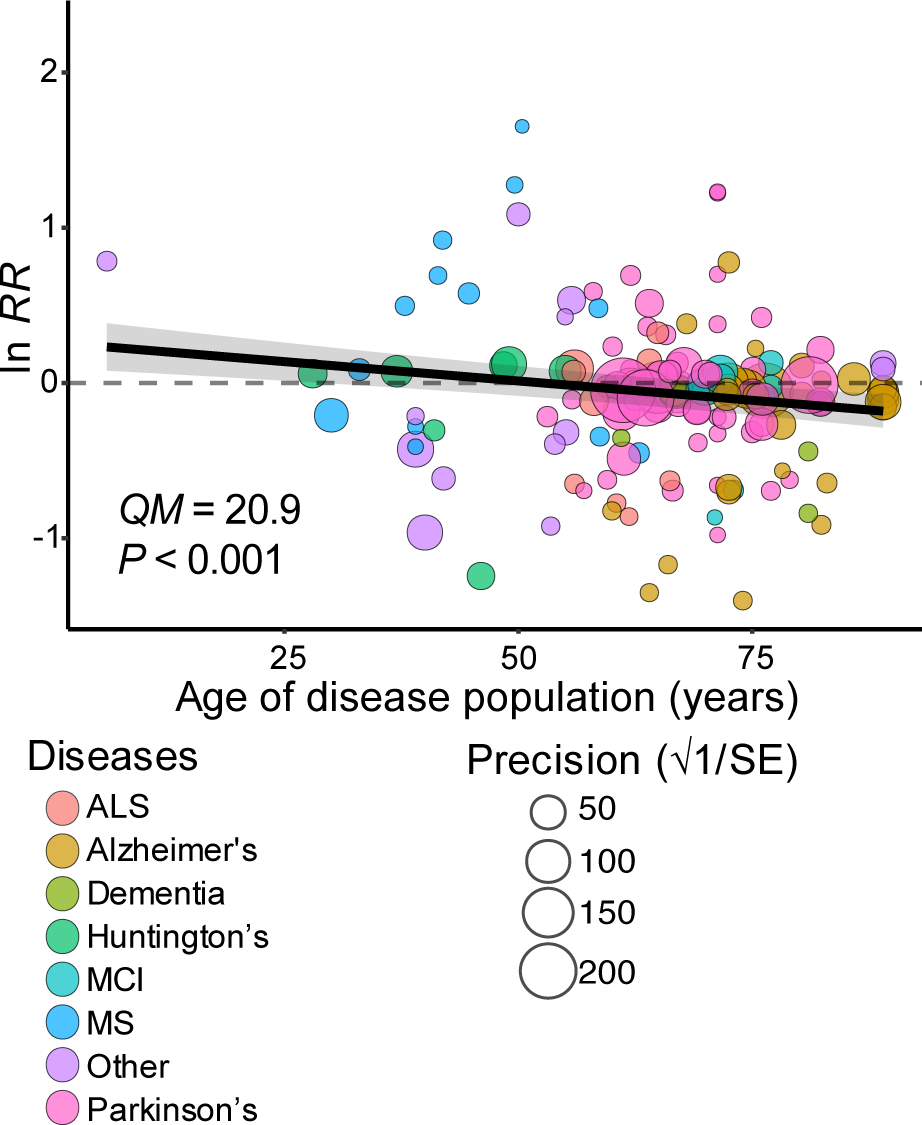
mtDNA-CN decreases more with neurodegeneration as age increases. Effect size is displayed on the y-axis (as on x-axis in Fig. 1), while age of the disease population is on the x-axis (see Fig. S3 for using age of control population). Points represent individual effect sizes scaled to precision (i.e., sample size) and colored by disease. *QM* and *P* statistics are displayed from a mixed-effects meta-analytical model with age of disease population as a sole moderator, with line of best fit and 95% CI also taken from this model.

Interestingly, the tissue used to measure mtDNA-CN did not explain a significant proportion of the heterogeneity in effect sizes (Fig. 2B, *P* = 0.600). The biggest difference was between mtDNA-CN measured in cerebrospinal fluid (13% ± 10% decrease in disease) vs. whole blood (3% ± 7% increase in disease), but this comparison was not statistically significant and both measures were not significantly different from no difference in mtDNA-CN between populations (*P* > 0.162).

Other moderators investigated did not explain a significant amount of heterogeneity in the effect sizes (Fig. 2, S2-S5, Table S4), including which location the population was from, how many or which mtDNA markers were used to estimate mtDNA-CN, and whether mtDNA-CN was normalized to a nuclear marker (*P* > 0.439 for all). When nuclear markers were used, the number or genomic region used did not explain a significant amount of heterogeneity in the effect sizes (*P* > 0.295 for both). The results also did not vary significantly based on whether the disease and control populations were matched for age or with the absolute difference in age between populations (Fig. S2, S5, *P* > 0.545 for both). Assay method had a stronger, but statistically non-significant effect on results (Fig. 2, *P* = 0.142), such that studies using the two most popular methods (qPCR and dPCR) had somewhat opposite results, with qPCR studies showing a 10% decrease and dPCR studies showing an 18% increase in mtDNA-CN with disease, but both methods overlapped a zero effect size. The author that extracted the data also had little effect on results (*P* = 0.615).

### Importance, interactions, and confounding of moderators

Using AIC values to further investigate significant moderators, we found that disease diagnosis explained the most variation in effect sizes, followed by sex, then age of the population (Table S5). There was a significant interaction between age of the population and disease type (*P* < 0.001), such that for most diseases mtDNA-CN decreased in disease populations as age increased, but for ALS (*P* = 0.072), MS (*P* = 0.108), and Parkinson’s (*P* = 0.231) mtDNA-CN tended to increase in disease populations compared to control ones as age increased (Fig. S6). There was also a significant interaction between age and sex of the population (*P* < 0.001), such that as populations got older, mtDNA-CN decreased significantly more in the disease population when both sexes were even (*P* < 0.001), but did not decrease significantly in male- or female-biased populations with age (*P* > 0.086 for both, Fig. S7). There was also a significant interaction between the age of the population and the tissue examined (*P* < 0.001), although mtDNA-CN decreased to at least some degree in the disease population as populations got older across most tissues (*P* < 0.046 for all, Fig. S8).

Interactions between categorical moderators were also statistically significant, but for many of these combinations there were a small number of effect sizes, or none at all, making inferences difficult. Therefore, here we only highlight a few interactions that might spur future research. For example, for Parkinson’s disease, effect sizes tended to be negative for male-biased populations, but were positive for the few female-biased populations, suggesting lower mtDNA-CN with Parkinson’s might be more extreme in males (Fig. S9). Brain tissue generally showed lower mtDNA-CN in disease populations across most diseases, but the cellular fraction of blood yielded different results for Alzheimer’s vs. Parkinson’s diseases (Fig. S10), suggesting that while tissue was not a significant predictor in single-moderator models, mtDNA-CN in some tissues might be better biomarkers for specific diseases. For example, low mtDNA-CN in cerebrospinal fluid may be a marker for Parkinson’s disease, while high mtDNA-CN in cerebrospinal fluid may be a marker for MS (Fig. S10).

We also investigated whether moderators were confounded with each other to better interpret results. For example, if all studies of ALS were female-biased, then it would be impossible to disentangle the effects of disease from sex. We did find significant confounding of several moderators, which influenced our analyses and should be considered when interpreting results. For example, disease diagnosis and disease neuropathology were nearly totally confounded (e.g., all Parkinson’s studies were categorized as synucleinopathies). We therefore focused only on disease diagnosis in most models. Similarly, average ages of disease and control populations were highly correlated (*P* < 0.001, *r*^2^ = 0.72, Fig. S11), prompting us to only consider age of the disease population in most analyses. While disease and sex were also statistically confounded (*P* < 0.001, chi-squared test), some diseases with larger numbers of effect sizes included studies with different sex ratios.

Similarly, a wide range of tissues were only investigated for diseases represented by large numbers of effect sizes, making inferences somewhat difficult. For example, all Huntington’s studies examined mtDNA-CN in the cellular fraction of blood. Sex and tissue were statistically confounded (*P* = 0.024), but many tissues were represented by populations with different sex ratios. Age was also different across diseases, tissues, and sex (*P* < 0.003 for all, Fig. S12-S14), which has important implications for interpreting our results. Alzheimer’s studies tended to focus on older populations, which may explain why mtDNA-CN was especially low in this disease if older populations show decreased mtDNA-CN with neurodegeneration. Similarly, female-biased populations tended to be older. Finally, as expected, brain tissue was from older individuals, which may explain bigger effects in this tissue in some analyses.

We next tested models with multiple moderators and their interactions to reduce heterogeneity as much as possible among effect sizes. While more complex models did reduce heterogeneity substantially (Table S6), even the most complex model still had excessive heterogeneity (*I*^2^ = 86%, *Q* = 193.6, *P* < 0.001). This most complex model included all biological moderators that were not heavily confounded and the interactions between them: disease diagnosis, age of the disease population, sex, tissue, assay method, and location.

### Effects become more moderate over time

We next investigated how effect sizes changed over time. When year of publication was used as a moderator by itself (as above for the other moderators), it explained a significant amount of heterogeneity in the effect sizes (*P* = 0.030, Table S4, Fig. 4A). As the year of publication became more recent, effect sizes became more positive. For example, for a study published in the year 2000 (the earliest study included here), our dataset predicts mtDNA-CN would be decreased by 39% in disease populations, but by the year 2025 (the most recent study included here), mtDNA-CN was predicted to be the same in disease and control populations.

**Fig. 4.**
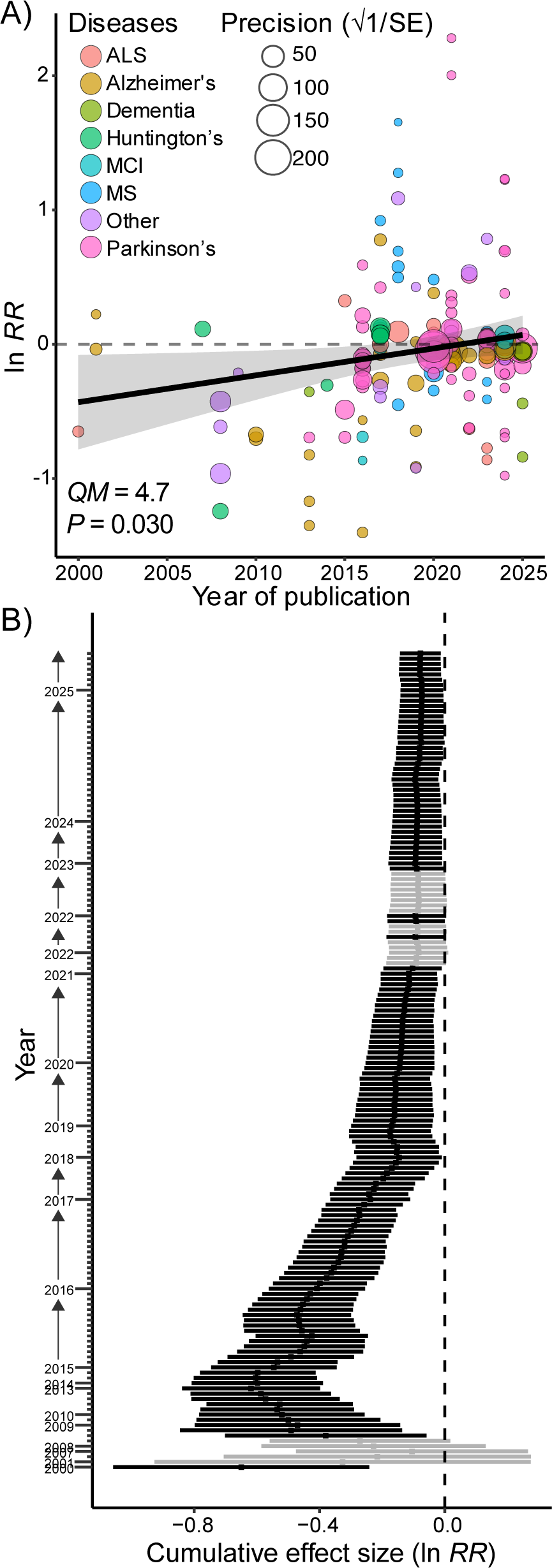
Earlier studies report more extreme decreases in mtDNA-CN with neurodegeneration. A) Year of publication explains significant heterogeneity in effect sizes (display as in Fig. 3). B) Results of cumulative meta-analysis, with sequential meta-analyses adding more effect sizes by year of publication and ascending on the y-axis. Points represent the grand mean effect size of each successive meta-analysis and lines represent 95% CIs (colors indicate whether an individual grand mean effect size range includes ln *RR* = 0).

We also performed a cumulative meta-analysis to investigate how effect sizes changed over time. In this technique, the earliest two effect sizes in the dataset are used to perform a meta-analysis and then sequential meta-analyses are performed by adding in effect sizes based on when they were published, with the last meta-analysis including all effect sizes (46). Supporting the above results, we found that earlier meta-analyses in this approach had significantly negative effect sizes, but later meta-analyses had more moderate effect sizes, suggesting earlier studies found lower mtDNA-CN in disease populations (Fig. 4B). For example, early overall effect sizes in the cumulative meta-analysis suggested mtDNA-CN was ∼85% lower in disease populations, but later ones put the estimate near 7% with borderline statistical significance.

### Little evidence of publication bias or sensitivity in effect sizes

As expected given the overall moderate effect sizes reported, we found limited evidence of publication bias. Funnel plots were largely symmetrical (Fig. S15), as confirmed by Egger’s regression analysis (*z* = 0.62, *P* = 0.535). Similarly, a trim-and-fill analysis suggested only 14 effect sizes were missing (Fig. S15). Importantly, these “missing” effect sizes were all for studies where mtDNA-CN would be lower in disease populations, suggesting the overall trend of lower mtDNA-CN with disease is not likely due to missing publications with the opposite effect. Fail-safe numbers were also high (*n* > 20,000), suggesting many thousands of effect sizes would need to have been missed in our search to offset the observed results. Leave 1 out analysis also suggests the overall estimated effect size was robust to excluding individual effect sizes, as repeating the meta-analysis after excluding any one effect size always yielded a similar overall effect size (ln *RR* range: −0.087 to −0.070). Influencer analysis yielded similar results, with only two effect sizes identified as significant influencers.

We also used a model with publication as a single moderator to confirm that effect sizes varied among publications. As expected, publication explained a large amount of heterogeneity in the effect sizes (*P* < 0.001, Table S4). Although not surprising, this further supported using publication as a random effect in the main meta-analysis models.

## Discussion

### Decreased mtDNA-CN is not a reliable biomarker for neurodegeneration

Our accumulated dataset from 76 publications included 156 instances where a population with a diagnosed neurodegenerative disease was compared to a control population, with 39,165 participants being examined across all studies. Despite gathering the largest such dataset to date, we did not find a consistent, statistically significant difference in mtDNA-CN between people with and without a diagnosed neurodegenerative disease. Although the overall grand mean effect size did trend towards lower mtDNA-CN with neurodegeneration, the effect was modest (only a ∼5% decrease) and statistically indistinguishable from no change after applying appropriate models. Heterogeneity was extreme in all meta-analytical models we investigated, suggesting effect sizes vary widely among individual studies. Recent analyses came to similar conclusions, with effects being modest, inconsistent, and not statistically robust (21). Even when confined to single diseases and tissues, patterns can be unclear or “controversial” (35).

Because mitochondria are central hubs for key metabolic processes, mitochondrial function is linked to many physiological processes. mtDNA-CN is accordingly influenced by myriad factors, which complicates detecting any underlying patterns between mtDNA-CN and neurodegeneration. For example, while low mtDNA-CN is associated with many diseases (13, 23, 51–53), an analysis of UK Biobank data from > 250,000 individuals found that mtDNA-CN in blood also varied depending on blood composition, time of day, month, and fasting condition (54). When these covariates were controlled for, mtDNA-CN was not lower in diseased individuals compared to control individuals, including for dementia (54). While many studies in our dataset did control for blood composition by limiting their tissue to specific types of blood cells (31, 55–59), controlling for nuances such as the time-of-day samples were collected was rarely reported. We also note that dementia was one of the few diseases where mtDNA-CN was significantly lower in our dataset (Fig. 1), which should be treated with caution given these additional complications and the UK Biobank results (54).

An additional complication is that tissues that are easily accessible in the clinic, such as different blood fractions, may not be a good proxy for the tissues most affected by neurodegeneration. MtDNA-CN can vary ∼50-fold across tissues in humans (60), but few studies have systematically measured mtDNA-CN simultaneously from multiple tissues across the same individuals, making it difficult to disentangle within- vs. among-individual variation in mtDNA. Surprisingly, the tissue examined did not explain a significant amount of heterogeneity in our dataset, with no single tissue showing a significant difference in mtDNA-CN between disease and control populations (Fig. 2B). However, we had to consolidate different tissues into the same category to generate enough power to compare among categories. For example, the cellular fraction of blood category in our dataset lumped together both leukocytes and platelets, which could show different or even opposite patterns in mtDNA-CN during neurodegeneration. Moreover, tissue did seem to be important in specific diseases. For Parkinson’s disease (the most studied disease in our dataset), cerebrospinal fluid did show lower mtDNA-CN during neurodegeneration, while other tissues did not (Fig. S10). For some diseases, mtDNA-CN was only quantified in a limited number of tissues, limiting inferences. For example, no studies of ALS quantified mtDNA-CN in cerebrospinal fluid (Fig. S10), which was was reported in a recent systematic review (21). Some neurodegenerative diseases may also affect specific brain regions or neuron types (61, 62), resulting in a natural target for where changes in mtDNA-CN should be most apparent if linked to disease. For Parkinson’s, the selective loss of dopamine neurons in the substantia nigra is a defining pathological feature, and studies that examined neurons from this region (56, 63, 64) did find a significant decrease in mtDNA-CN, while other brain regions showed no change (*P* < 0.001, Fig. S16).

### How disease, age, sex, and other factors influence mtDNA-CN during neurodegeneration

A recent systematic review stated that the poor reproducibility in results among neurodegeneration studies investigating mtDNA-CN is “likely due to the lack of consideration of the many factors known to affect mtDNA levels” (21). A main goal of our study was to investigate potential moderators to explain this variation among effect sizes. Importantly, many of the moderators we considered important *a priori* did not explain a statistically significant amount of variation (Fig. 2, Fig. S2-S5). Which tissue was examined did not affect overall results (see above), along with other methodological considerations such as which or how many mtDNA or nuclear DNA markers were used or the assay method. We expected the region of mtDNA to be important because certain stretches of mtDNA are more likely to undergo large deletions than others (65, 66). While surprising, this does offer some practical benefits for quantifying mtDNA-CN in disease vs. control populations. Sequencing approaches allow more powerful ways to quantify mtDNA-CN and can also be used to estimate heteroplasmy or the fraction of deletion-bearing mtDNA genomes (67, 68) and digital PCR approaches are more precise than qPCR (69), the more traditional method of quantifying mtDNA-CN. However, sequencing-based approaches and dPCR tend to be more expensive per sample and require more specialized equipment unavailable to most labs. qPCR produced comparable results to these more powerful methods (Fig. 2C), suggesting this traditional assay can continue to be relied on. Moreover, whether studies normalized mtDNA-CN to a nuclear control marker or not did not significantly influence results (Fig. S2). Although *absolute* quantification of the number of mtDNA genomes per cell requires some method of knowing how many cells are in each sample, this result suggests that estimating *relative* mtDNA-CN between disease and control samples may be accomplished without normalization, assuming sample collection methods are the same across populations. Normalizing to a nuclear marker or the number of cells examined may also be unnecessary for some tissues that are multinucleated or when examining cell-free DNA.

Although most methodological considerations were not major moderators of note, biological differences tended to explain more heterogeneity among studies. There were notable differences depending on which neurodegenerative disease was examined, with Alzheimer’s showing a convincing 21% decrease in mtDNA-CN compared to controls (Fig. 1). Other diseases such as Huntington’s and MCI also showed a statistically significant decrease in mtDNA-CN, but were based on fewer studies. No disease showed a statistically significant increase in mtDNA-CN, although MS came closest (17% increase ± 31%) as suggested when examining cerebrospinal fluid previously (21, 70–72). This finding suggests that mtDNA-CN may be an especially useful biomarker for specific neurodegenerative diseases such as Alzheimer’s, but other well-studied diseases in our dataset such as Parkinson’s may be more complicated. Populations with Alzheimer’s also showed the most extreme depletion of mtDNA in brain tissue (∼60%), which suggests that mtDNA-CN depletion may be biologically relevant for Alzheimer’s progression or severity. As mentioned above, specific tissues may be especially useful in some diseases, so even when overall effect sizes were moderate in our meta-analyses, mtDNA-CN should not be abandoned as a potential biomarker.

Disease explained more heterogeneity than any other single moderator in our dataset (Table S4-S5), but sex and age were also significant (Fig. 2, 3). However, overall effects of sex may be an artifact of the small number of studies that used populations biased towards males or females, as both male- and female-biased populations showed increased mtDNA-CN with neurodegeneration compared to populations with relatively even proportions of males and females (Fig. 2A). If this was a true biological effect, we would expect males to show one pattern, females to show the opposite, and matched populations to show an intermediate pattern. This is largely what was seen for Parkinson’s (Fig. S9), one of the few diseases with data available for male-biased, female-biased, and sex-matched populations. In Parkinson’s, males showed an extreme decrease in mtDNA-CN with neurodegeneration, females showed an increase, and matched populations were intermediate. This makes some biological sense, given an increased risk of developing Parkinson’s in males and other sex differences in disease progression, symptoms, and treatment response (73). However, studies of mtDNA-CN in female-biased Parkinson’s populations were rare and had small sample sizes (39, 74).

Age was a more convincing biological moderator of mtDNA-CN during neurodegeneration (Fig. 3, S5-S8). Across our analyses, older populations saw a more pronounced decrease in mtDNA-CN with neurodegeneration compared to younger populations. This makes intuitive sense because mtDNA-CN naturally decreases with age in humans and other vertebrates (54, 75, 76) and neurodegeneration may exacerbate this process. Moreover, most neurodegenerative diseases in our dataset are more pronounced at older ages. Some studies examined mtDNA-CN in disease and control populations across wide age ranges and reported data as a function of age, which would allow for explicitly calculating an interaction between age and disease status within a study (32). However, data were rarely presented in this fashion.

Despite disease, age, and sex explaining a significant amount of heterogeneity in effect sizes, caution should be used when interpreting these results due to significant interactions and confounding among these moderators. For example, while mtDNA tended to decrease with neurodegeneration at older ages, within a given disease the effect was not very noticeable, even for diseases like Parkinson’s and Alzheimer’s with large sample sizes (Fig. S6). Population ages were also not evenly distributed across diseases (Fig. S13), and studies investigating Alzheimer’s disease tended to especially focus on older populations. Because old age is confounded with Alzheimer’s disease in our dataset, it is difficult to conclude whether effect sizes are larger in Alzheimer’s disease, older populations, or both.

Although the moderators we investigated did explain some variation across studies, extreme heterogeneity in effect sizes remained even in our most complex meta-analytical models. When considering all moderators and their interactions, we still recovered *I*^2^ at 86% of a possible 100%, meaning that 86% of the variance among effect sizes is likely due to unexplained sources of variation (i.e., moderators not accounted for), not random sampling error. Although daunting, our study did span many diseases and tissues and a more focused meta-analysis would be expected to show lower heterogeneity. However, a recent meta-analysis focused solely on Parkinson’s disease also found a high *I*^2^ (93%) despite only including 20 effect sizes from 13 studies (35). Taken together, how mtDNA-CN changes with neurodegeneration appears very inconsistent across studies and identifying additional sources of this variation remains a challenge for future studies.

### Legacy effects may dominate perceptions of mtDNA-CN during neurodegeneration

MtDNA-CN has been associated with disease for 35 years, with original studies documenting severe mtDNA depletion in patients with known mitochondrial dieases (75, 76). Later, mtDNA levels were associated with chronic diseases such as cancer, obesity, and neurodegeneration (77), with more recent studies associating lower mtDNA levels with everything from psoriasis (78) to asthma (79). Although increased mtDNA levels can be associated with disease (most notably cancer), increased mtDNA seems to be most associated with decreased disease severity (80, 81). This has led to a general perception in the literature that when it comes to copies of the mtDNA, “the more the better” (24).

For neurodegeneration, early studies that found lower mtDNA in disease populations may have disproportionately shaped the field. We found that effect sizes became more moderate in later years, with cumulative meta-analysis showing overall negative effect sizes until the early 2020s (Fig. 4). This could be due to biological effects, as the earliest studies in our dataset focus on diseases other than Parkinson’s (50, 82, 83). It could also be due to changing methodologies, although qPCR use dominated both earlier and later studies. However, we favor the interpretation that earlier studies may have overestimated effect sizes because they relied on smaller sample sizes. The average sample size for the first 20 effect sizes in our dataset was 36 disease individuals, while the average for the last 20 effect sizes was 103. Previous work has shown that the lower the sample size, the more likely an individual study is to overestimate an effect size (84, 85) and overall effects of interventions in pregnancy/perinatal and myocardial infarction studies tended to fluctuate widely until a large number of cumulative patients had been examined (86). Although publication bias was largely absent in our study, time-lag bias, where studies with smaller or statistically non-significant effects take longer to publish than those with larger effects, may also explain why effect sizes become more moderate over time in clinical medicine (87). In any case, finding extreme effects in earlier studies that become more modest after including later studies with more replicates is a somewhat common pattern in meta-analyses (88). We suggest the pattern observed here for mtDNA levels with neurodegeneration is due to a combination of earlier studies targeting systems where effects are particularly large and being based on smaller sample sizes. It is important that these “legacy” effects should not overshadow recent, more highly replicated studies.

### Missing studies, future directions, and promise for mtDNA-CN as a health biomarker

Several types of studies were missing or underrepresented in our dataset. As with many other clinical datasets, studies included here were largely from Europe, the United States, the United Kingdom, and East Asia (Fig. 2D). Only single studies were available from Africa and Australia and only a few studies were available from South/Central America and the Middle East (Fig. 2D). Future studies should focus on these underrepresented areas, as genetic makeup can strongly influence baseline mtDNA levels (54) and may also influence how mtDNA changes with neurodegeneration. Similarly, underrepresented diseases in our dataset such as dementia, Huntington’s, and MCI should be further examined to support the negative relationship between mtDNA-CN and neurodegeneration suggested here for these diseases based on a limited number of studies.

The number of effect sizes available was generally limited when considering combinations of specific diseases and tissues, ages, or other moderators (Fig. S6-S10). For example, it is impossible to assess how mtDNA-CN may change across tissues for Huntington’s disease because all studies included here focused on the cellular fraction of blood (Fig. S10). New studies explicitly focused on these missing combinations would be useful, but also very expensive and intensive. We offer two alternative suggestions. First, because we found age and sex to be important moderators across the overall dataset, it would be useful to study the effects of age and sex within an individual study. While these data likely exist, they are seldom reported in a usable way (although see (89) for an exception). Reporting these data would allow for detailed examination of how age and sex affect mtDNA-CN within specific diseases. Second, large biobanks and repositories of whole-genome sequence data across thousands of individuals provide an untapped resource for comparing mtDNA-CN across diseases, ages, and sex (54). These repositories often offer much larger sample sizes than those from individual, targeted clinical studies, even when not focused on populations that would be enhanced for chronic diseases (e.g., those recruited through a hospital). Finally, Biobanks are now available from areas of the world that were underrepresented in our dataset (90).

While the focus of our meta-analysis was on how and if mtDNA-CN changes with neurodegeneration, many of the studies included here also examined other effects or performed more thorough and nuanced correlations with mtDNA-CN. For example, Jędrak et al. (89) quantified mtDNA-CN in both symptomatic and pre-symptomatic populations with Huntington’s disease, finding elevated mtDNA levels in both groups when compared to controls. Similarly, studies in our dataset compared mtDNA levels before and after affected individuals were diagnosed (91), during disease progression associated with risk of developing dementia (92), and in early-onset vs. idiopathic Parkinson’s (93). Although we focused on the simplest comparison of control vs. disease populations with as few confounding complications as possible, further analyses of these trends in individual studies and future meta-analyses will be important in developing mtDNA levels as a biomarker for both general neurodegeneration and specific diseases.

Neurodegenerative diseases were the focus of our meta-analysis, but altered mtDNA levels have been associated with many other diseases and metrics of health (11–14). Although these associations are often based on studies with large sample sizes (94, 95) and narrative reviews are common (24, 96), formal meta-analyses are lacking in this field. Quantifying mtDNA levels in easily accessible tissues holds great promise as a general biomarker for health (12). However, inferences from single studies are often limited to specific populations and methods. Meta-analyses not only synthesize findings within a given field using appropriate statistical analyses, but also enable comparisons of factors that vary across studies and would be difficult to examine within a single study (97). We argue that additional meta-analyses of mtDNA levels in human health and disease are an important part of realizing the full potential of mtDNA as a biomarker.

## Acknowledgements

We thank members of the Havird and Donegan labs, participants in a recent meta-analysis graduate class at UT, and anonymous reviewers for comments that strengthened our study.

