## Supplementary material for "Mitochondrial DNA copy number in neurodegenerative diseases: a global meta-analysis of 156 comparisons across 76 studies": Fig. S

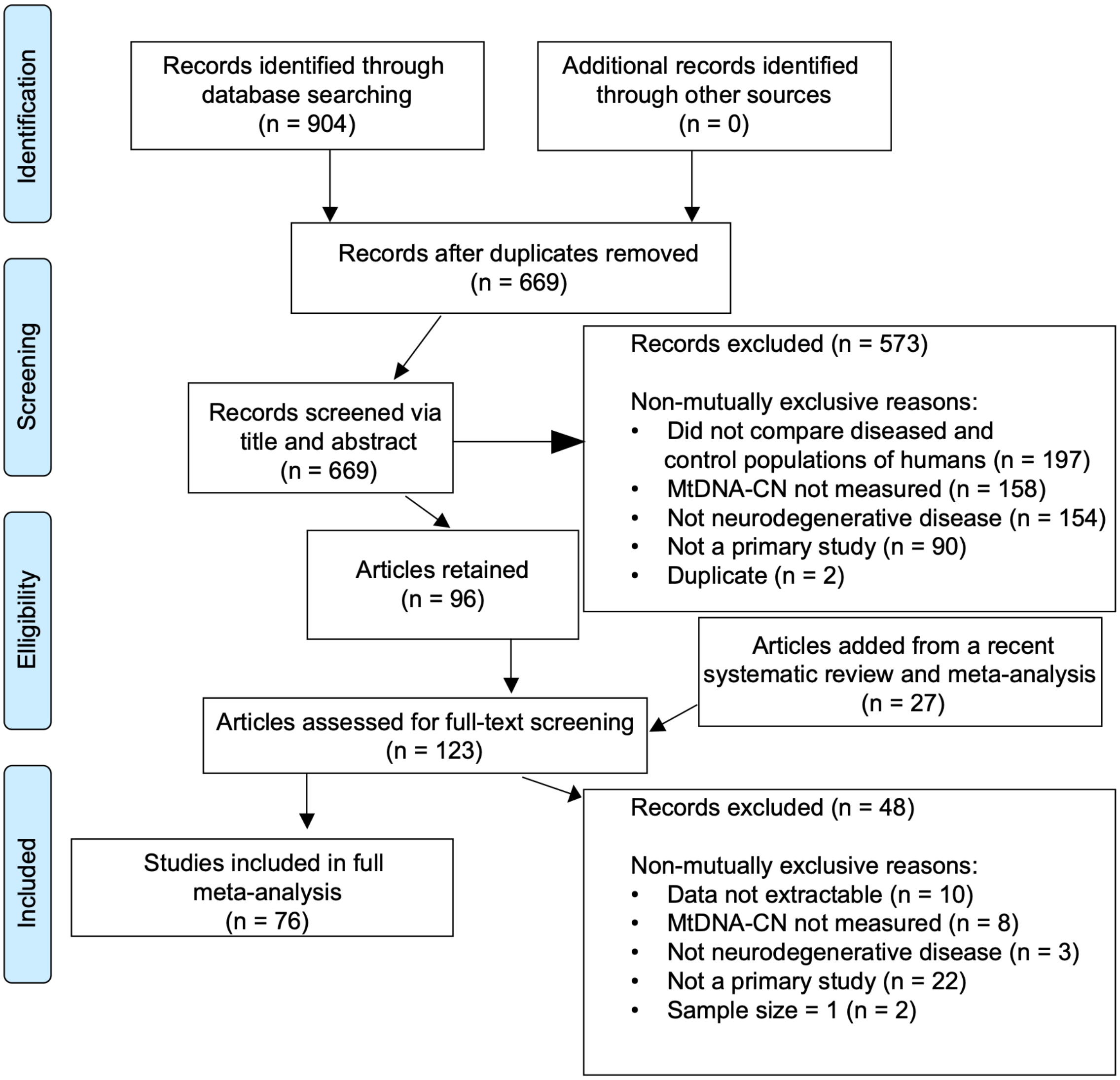


*From:* Moher D, Liberati A, Tetzlaff J, Altman DG, The PRISMA Group (2009). *P*referred *R*eporting *I*tems for *S*ystematic Reviews and *M*eta-*A*nalyses: The PRISMA Statement. PLoS Med 6(7): e1000097. doi:10.1371/journal.pmed1000097

**Fig. S1.** PRISMA-style diagram detailing how many records were examined from a systematic literature search, how many were excluded during different rounds of screening, and how many were ultimately included in the meta-analysis.

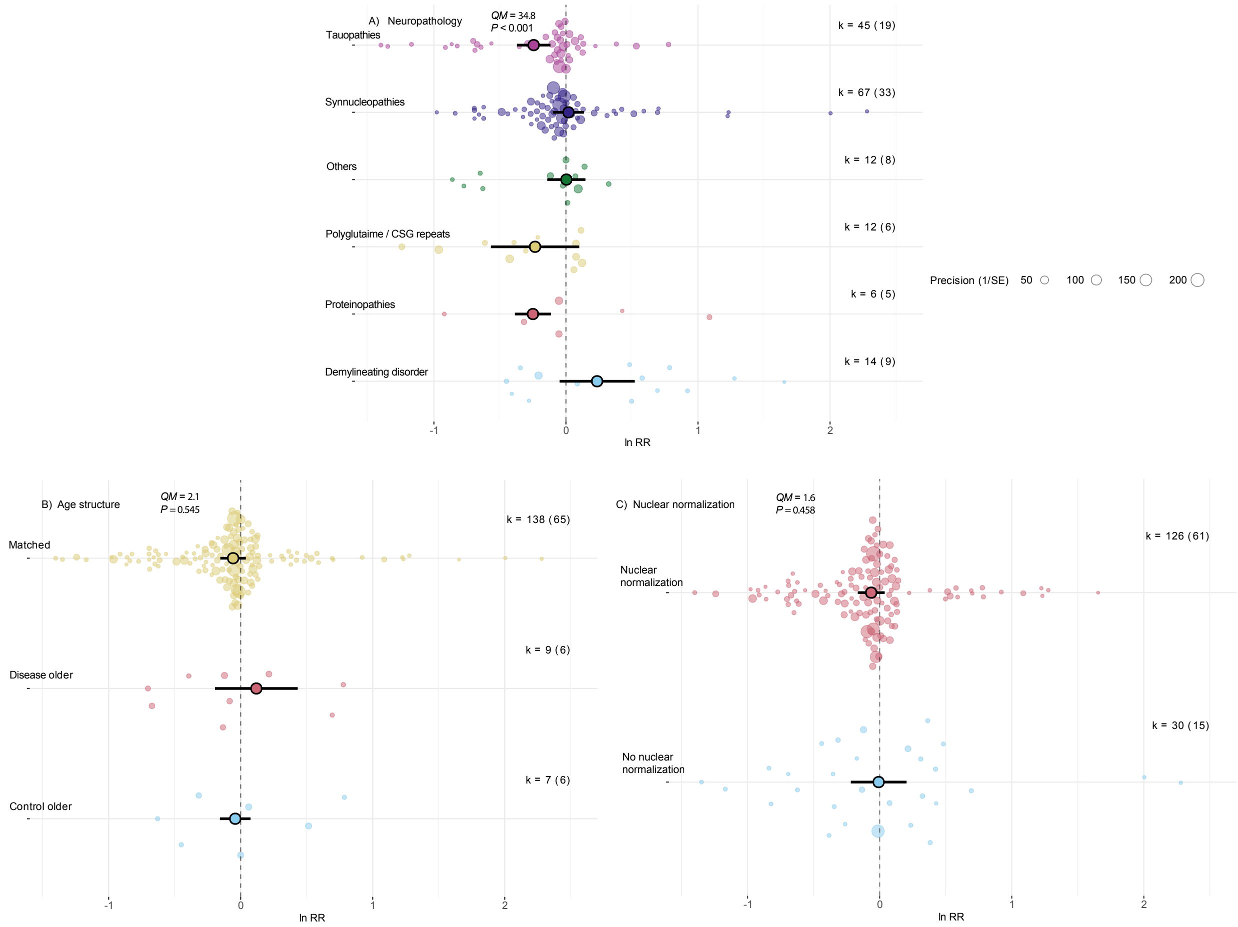


**Fig. S2.** Some factors explain why mtDNA-CN may or may not change with neurodegeneration. A) The neuropathology of the underlying disease explained significant heterogeneity among effect sizes, but the B) age matching structure of the disease and control populations and C) whether a nuclear marker was used to normalize mtDNA-CN did not. Effect sizes are displayed on the x-axes as in Fig. 1, along with *k* values. Transparent points represent individual effect sizes and are scaled to their precision (i.e., sample sizes), while opaque point represent mean effect sizes per category and are shown with 95% CIs. *QM* and *P* statistics are displayed from mixed-effects meta-analysis models using single moderators.

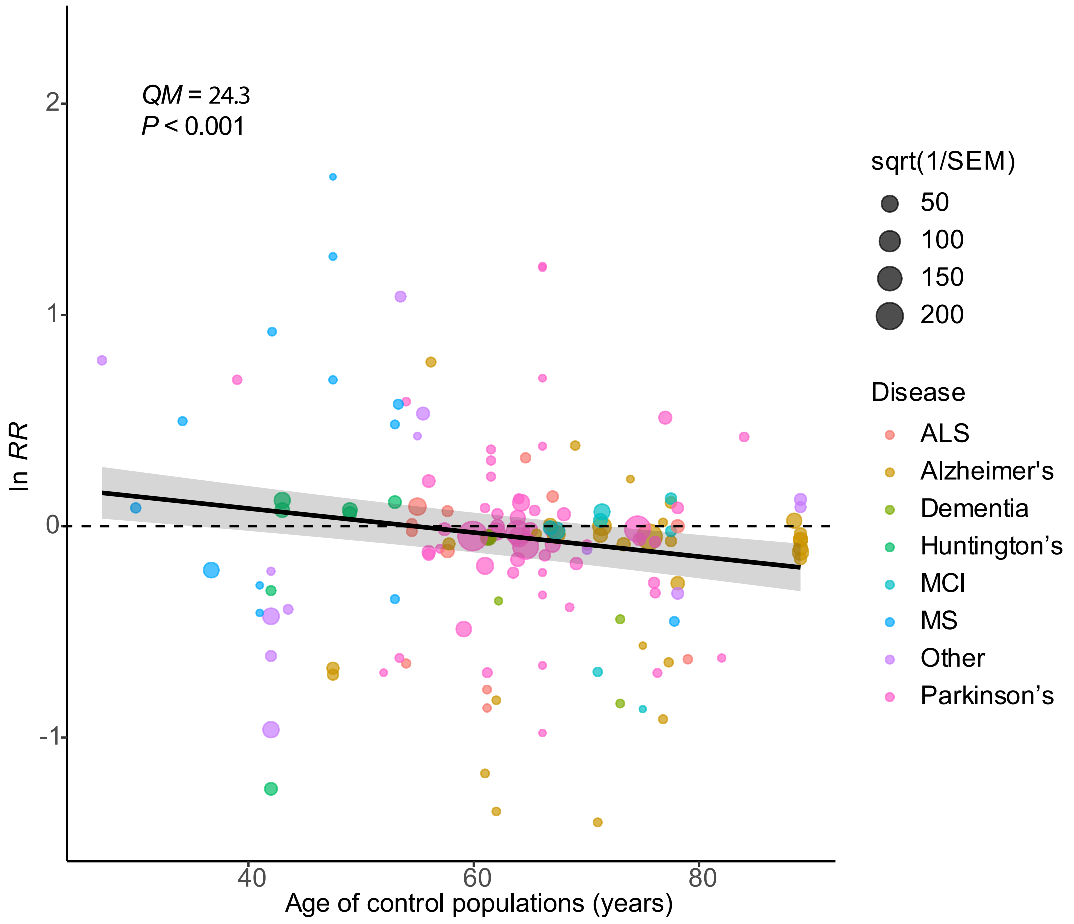


**Fig. S3.** mtDNA-CN decreases more with neurodegeneration as age increases. Effect size is displayed on the y-axis (as on x-axis in Fig. 1), while age of the control population is on the x-axis (see Fig. 3 for using age of disease population). Points represent individual effect sizes scaled to precision (i.e., sample size) and colored by disease. *QM* and *P* statistics are displayed from a mixed-effects meta-analysis model with age of control population as a sole moderator, with line of best fit and 95% CI also taken from this model.

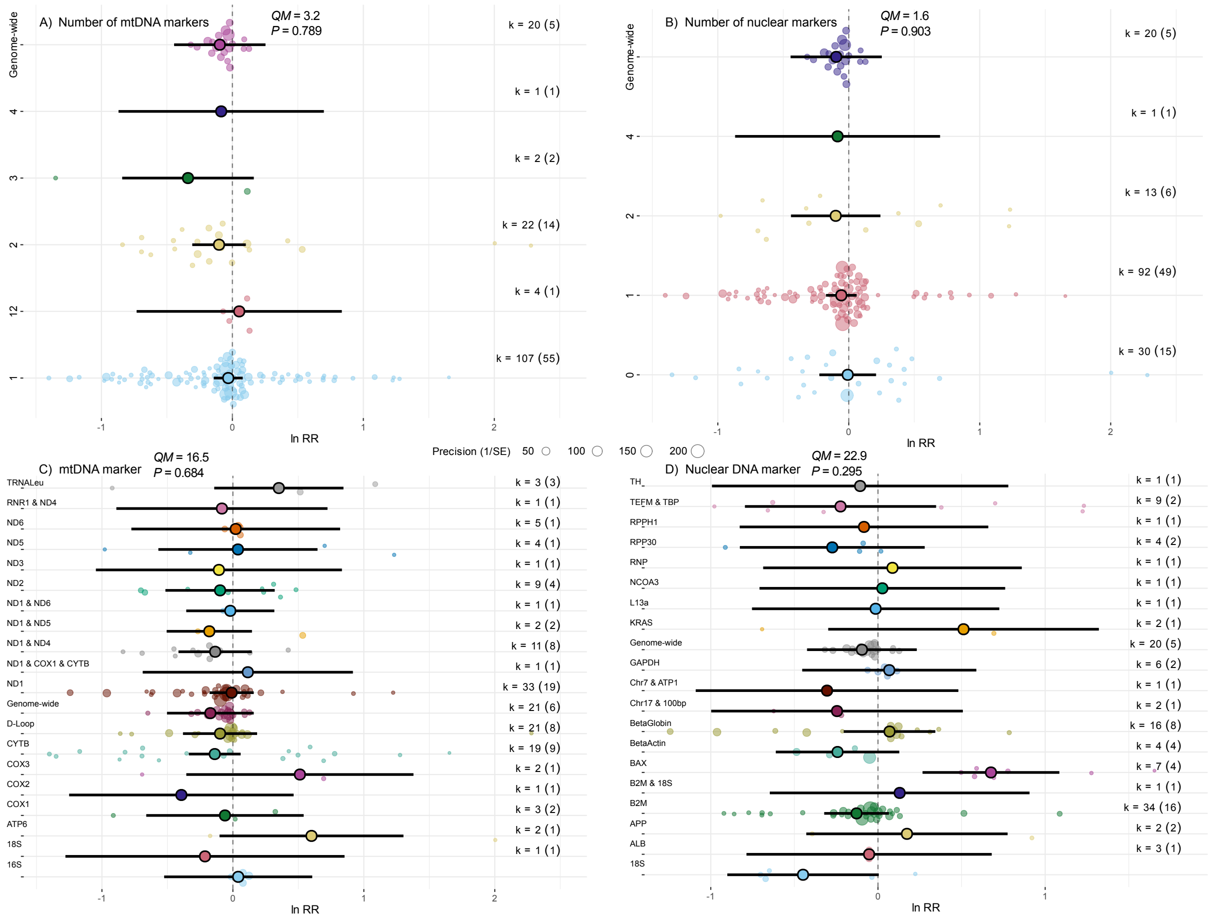


**Fig. S4.** Numbers and types of mtDNA and nuclear DNA markers do not explain why mtDNA-CN may or may not change with neurodegeneration. A) number of mtDNA markers used, B) number of nuclear DNA markers used, C) the mtDNA marker used, and D) the nuclear DNA marker used. Effect sizes are displayed on the x-axes as in Fig. 1, along with *k* values. Transparent points represent individual effect sizes and are scaled to their precision (i.e., sample sizes), while opaque point represent mean effect sizes per category and are shown with 95% CIs. *QM* and *P* statistics are displayed from mixed-effects meta-analysis models using single moderators.

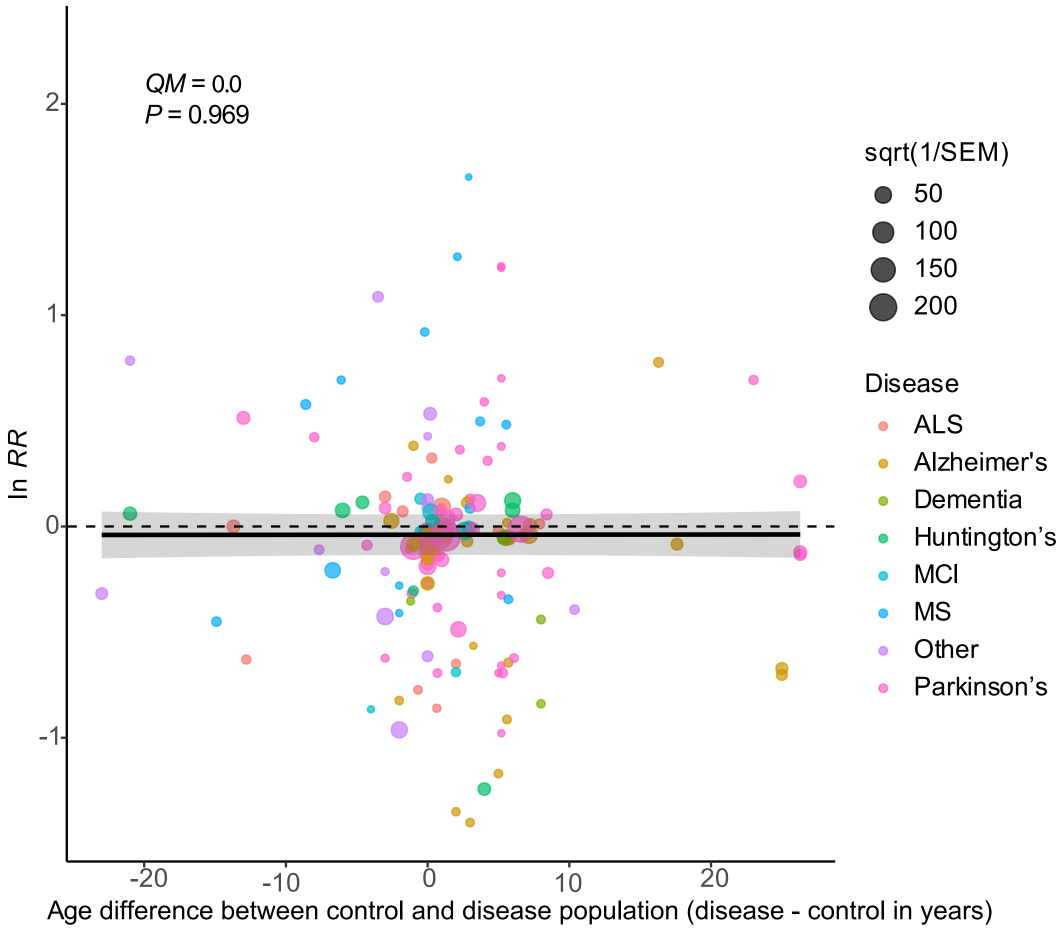


**Fig. S5.** mtDNA-CN does not change more or less with neurodegeneration as difference in average ages between the control and disease population changes. Effect size is displayed on the y-axis (as on x-axis in Fig. 1), while the age difference between populations (calculated as average age of disease – average age of control population) is on the x-axis. Points represent individual effect sizes scaled to precision (i.e., sample size) and colored by disease. *QM* and *P* statistics are displayed from a mixed-effects meta-analysis model with age difference as a sole moderator, with line of best fit and 95% CI also taken from this model.

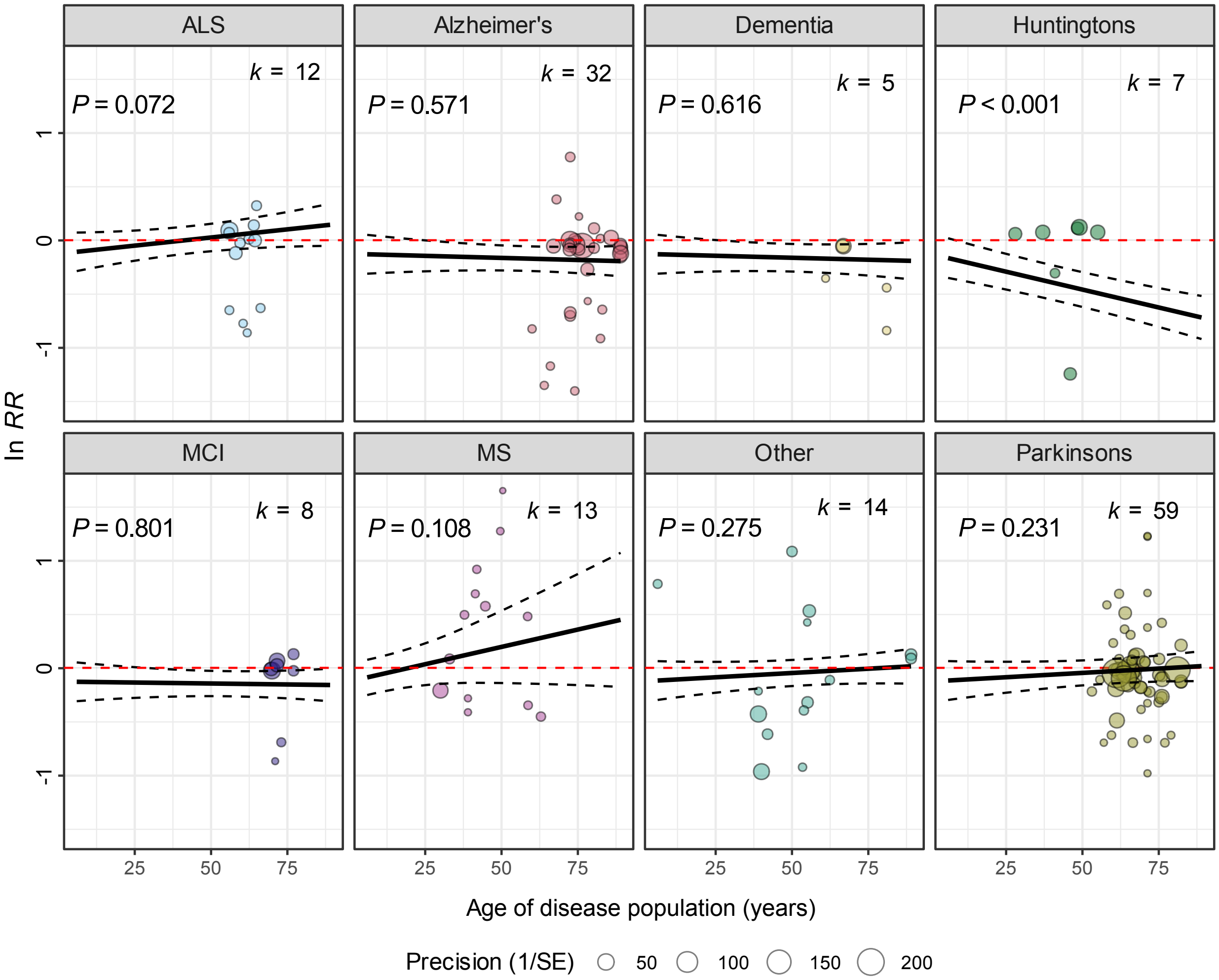


**Fig. S6.** How mtDNA-CN changes with neurodegeneration differs across age in different ways for different diseases. Effect size is displayed on the y-axis (as on x-axis in Fig. 1), while the age of the disease population is on the x-axis. Points represent individual effect sizes scaled to precision (i.e., sample size) and displayed and colored by disease. *QM* and *P* statistics are displayed from a mixed-effects meta-analysis model with the interaction between age of disease population and disease type as the sole moderator, with line of best fit and 95% CI also taken from this model.

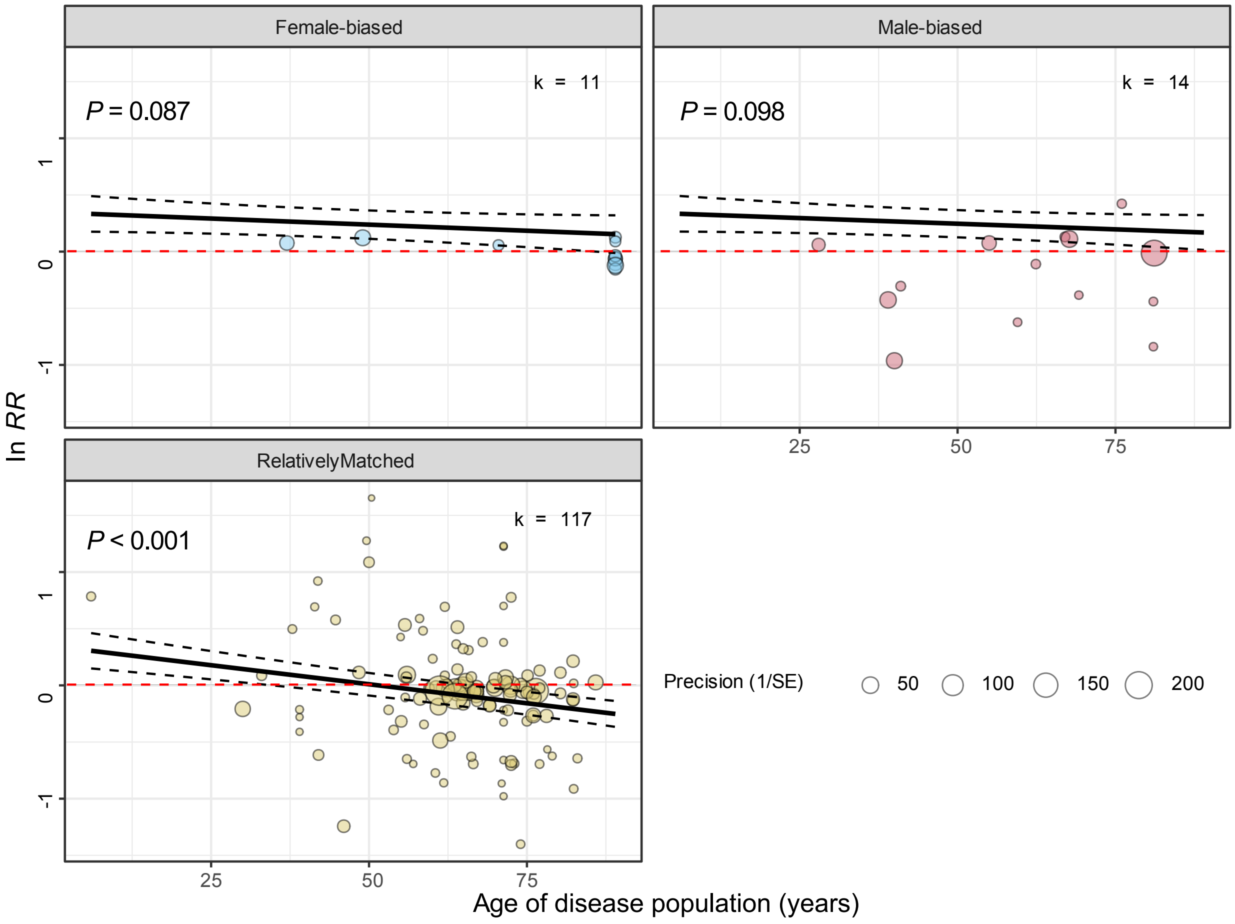


**Fig. S7.** How mtDNA-CN changes with neurodegeneration across age in different sexes. Effect size is displayed on the y-axis (as on x-axis in Fig. 1), while the age of the disease population is on the x-axis. Points represent individual effect sizes scaled to precision (i.e., sample size) and displayed and colored by sex ratio. *QM* and *P* statistics are displayed from a mixed-effects meta-analysis model with the interaction between age of disease population and sex ratio type as the sole moderator, with line of best fit and 95% CI also taken from this model.

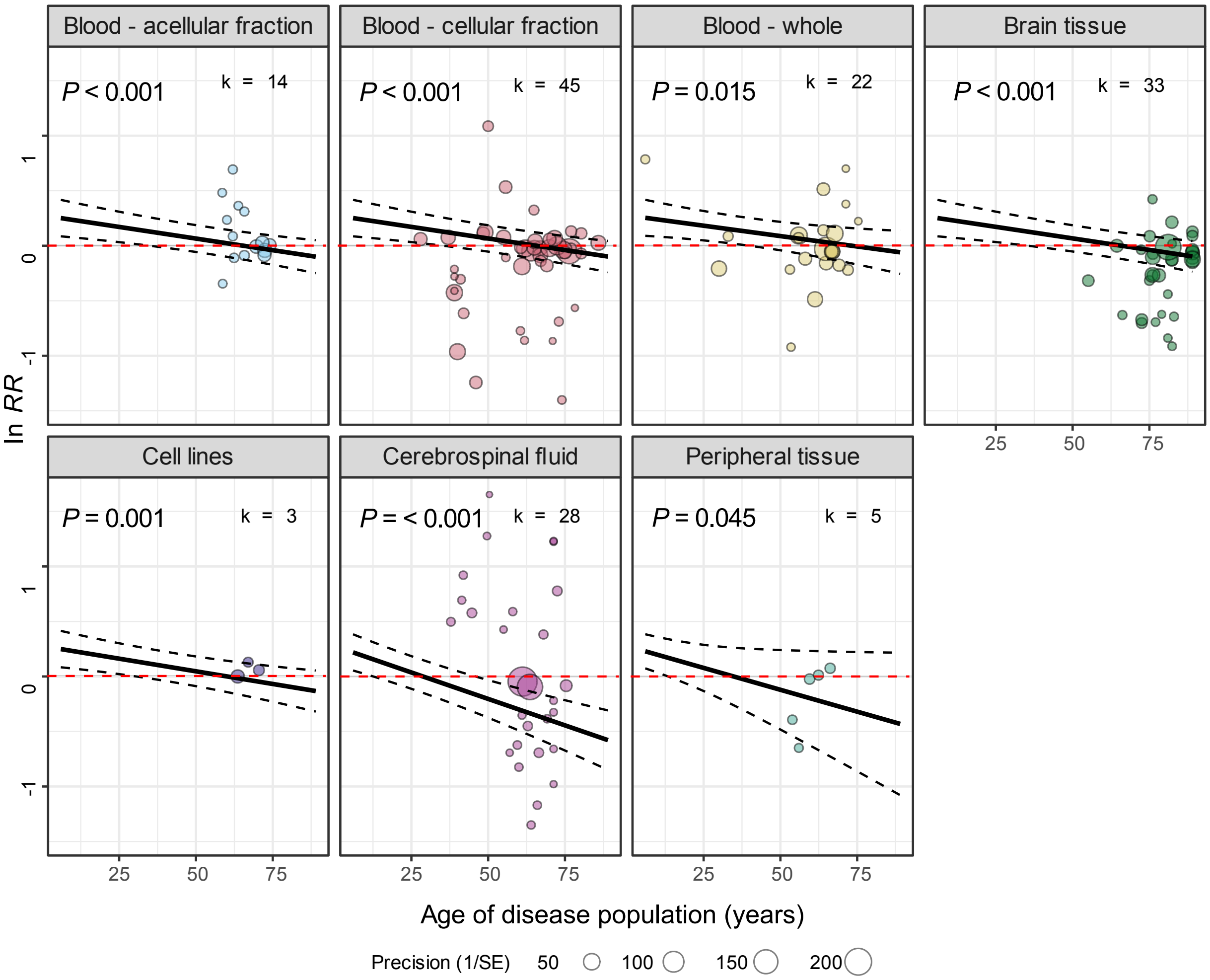


**Fig. S8.** How mtDNA-CN changes with neurodegeneration across age in different tissues. Effect size is displayed on the y-axis (as on x-axis in Fig. 1), while the age of the disease population is on the x-axis. Points represent individual effect sizes scaled to precision (i.e., sample size) and displayed and colored by tissue examined. *QM* and *P* statistics are displayed from a mixed-effects meta-analysis model with the interaction between age of disease population and tissue as the sole moderator, with line of best fit and 95% CI also taken from this model.

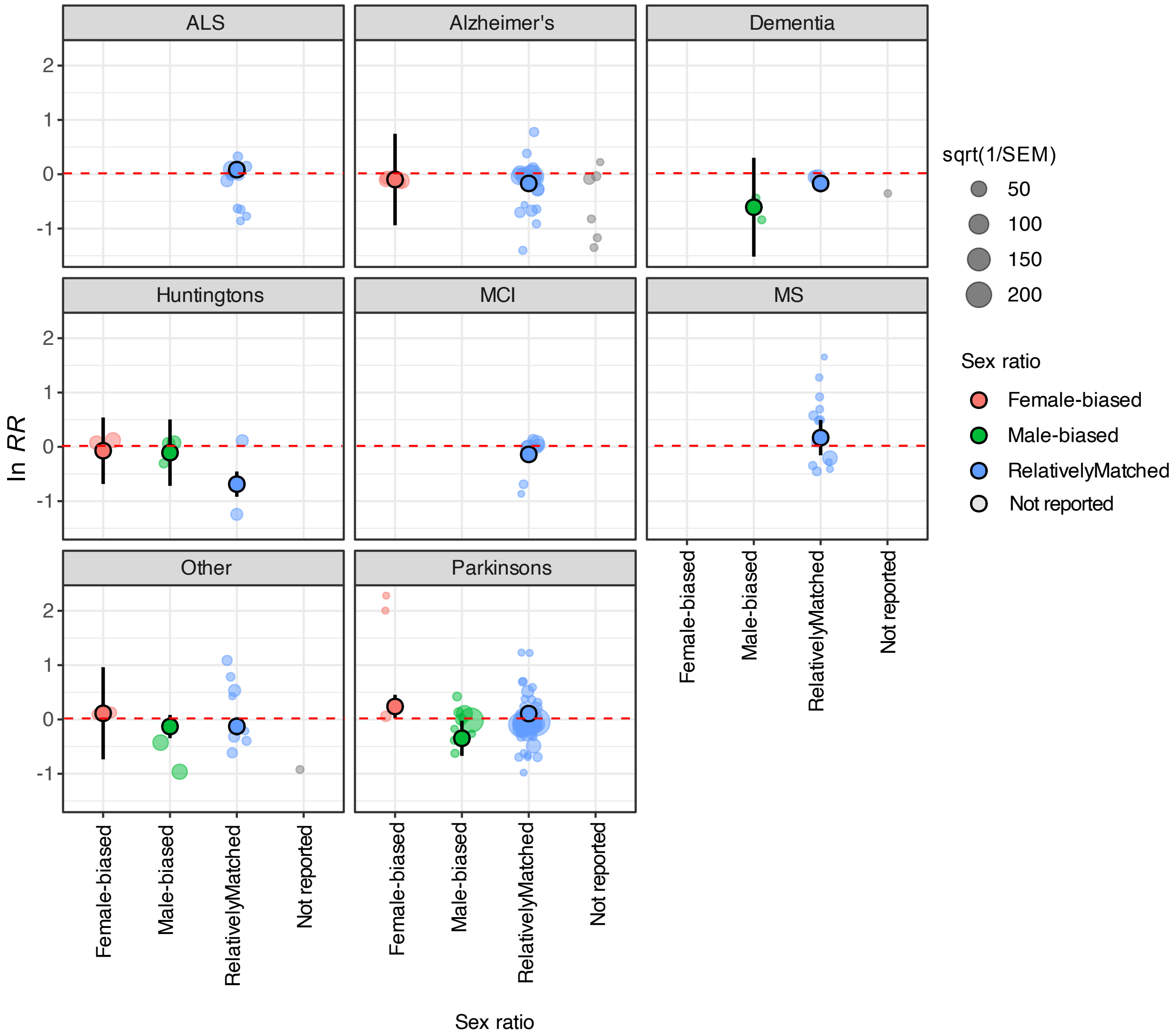


**Fig. S9.** How mtDNA-CN changes with neurodegeneration in different combinations of disease and sex ratio. Effect size is displayed on the y-axis (as on x-axis in Fig. 1), while the sex ratio is on the x-axis. Transparent points represent individual effect sizes scaled to precision (i.e., sample size) and displayed and colored by disease and sex ratio. Opaque points are mean effect sizes (with 95% CIs) for each combination of sex ratio and disease from a mixed-effects meta-analysis model with the interaction between sex ratio and disease as the sole moderator.

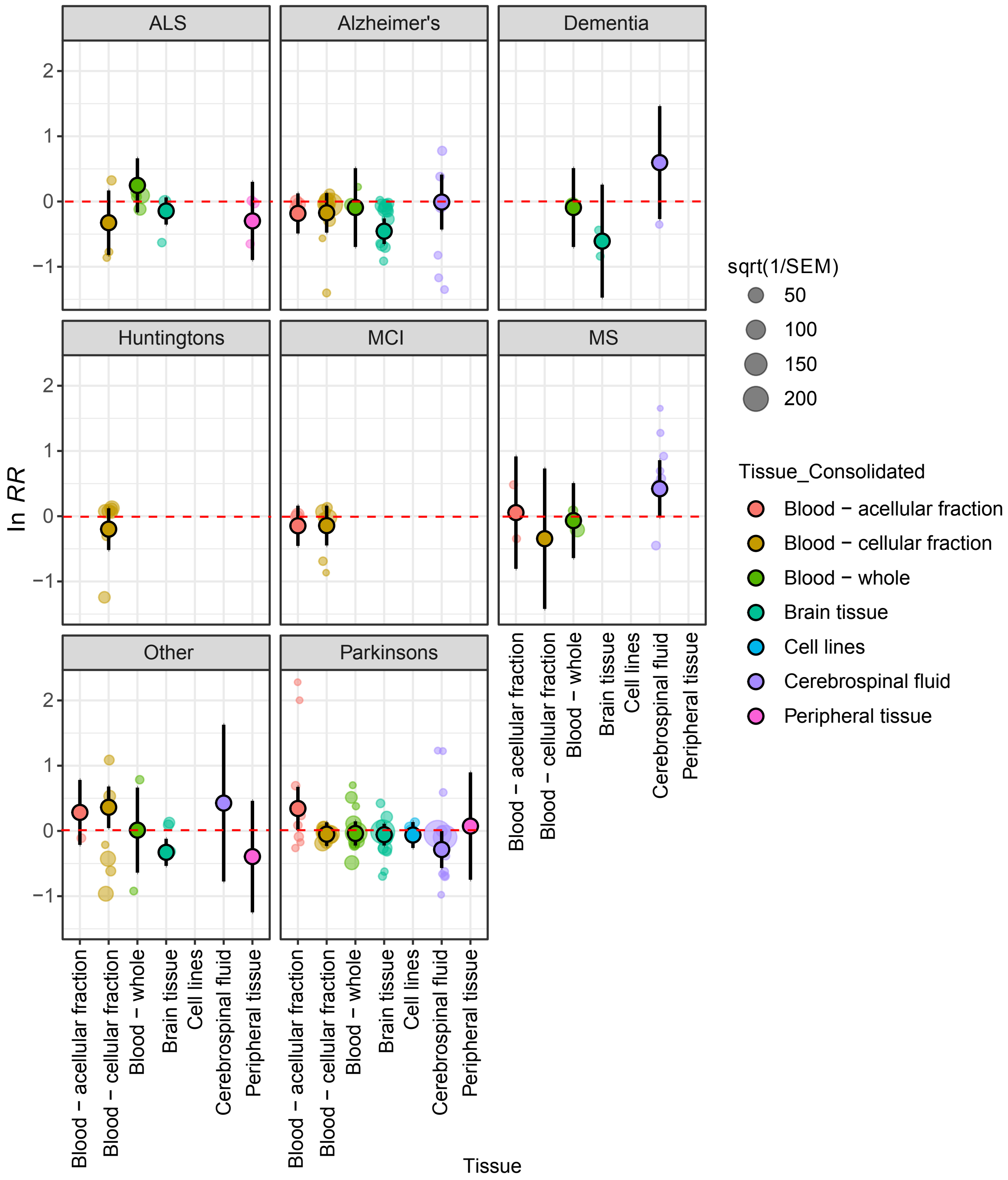


**Fig. S10.** How mtDNA-CN changes with neurodegeneration in different combinations of disease and tissue. Effect size is displayed on the y-axis (as on x-axis in Fig. 1), while the tissue is on the x-axis. Transparent points represent individual effect sizes scaled to precision (i.e., sample size) and displayed and colored by disease and tissue. Opaque points are mean effect sizes (with

95% CIs) for each combination of tissue and disease from a mixed-effects meta-analysis model with the interaction between tissue and disease as the sole moderator.

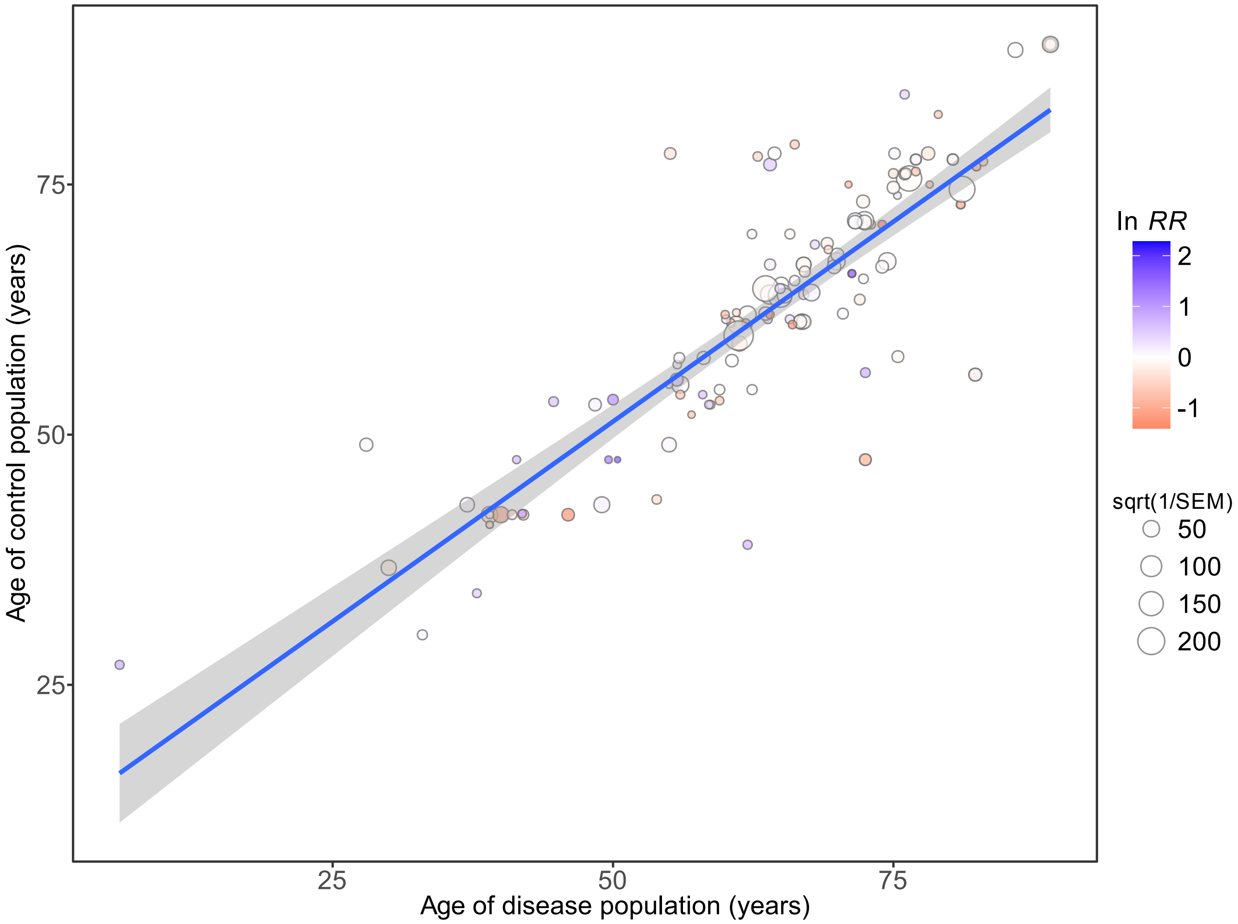


**Fig. S11.** Ages of disease and control population are strongly correlated in our dataset. Points are scaled by precision (i.e., sample size) and colored according to their effect size. Line of best fit and 95% CI are from a linear correlation model ($r^2$ = 0.72, $P$ < 0.001).

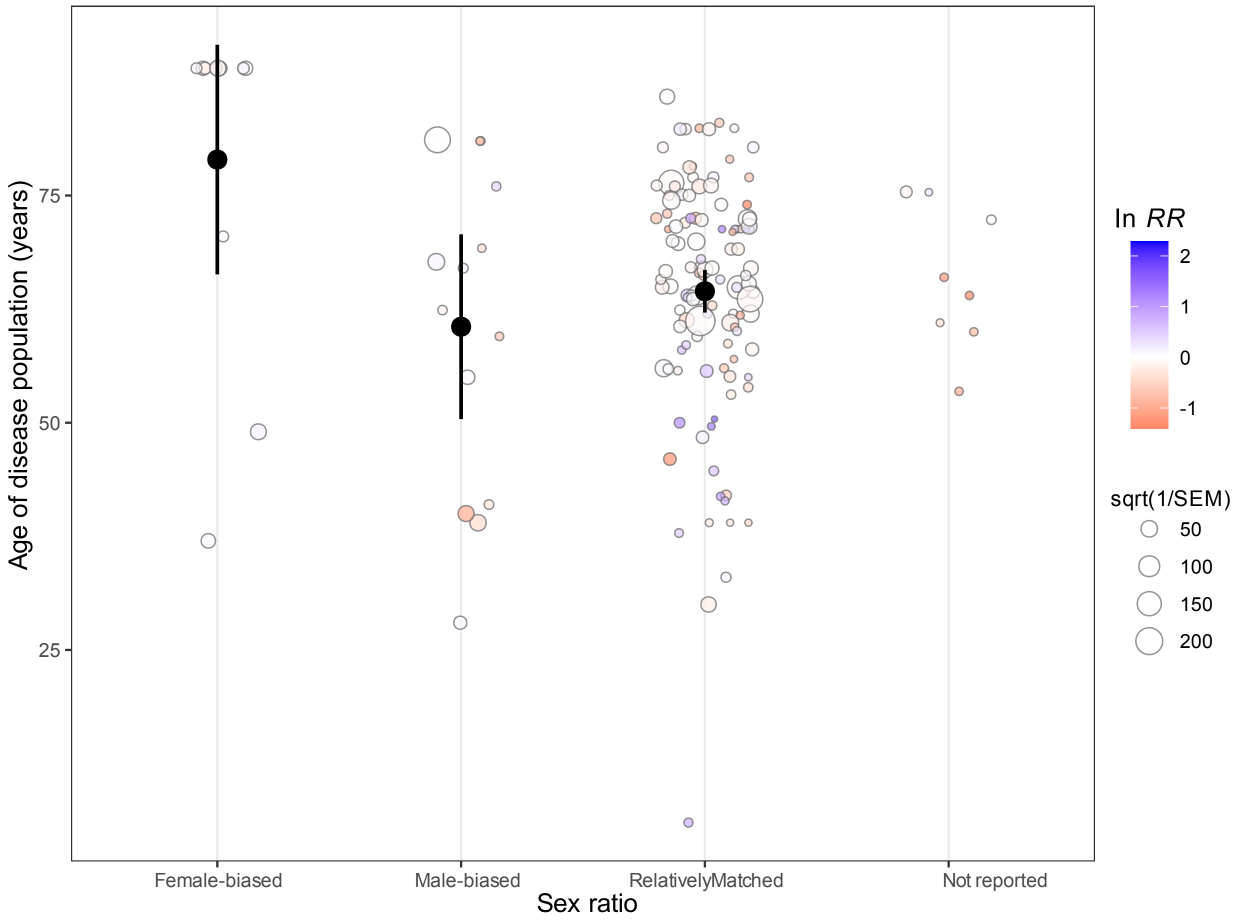


**Fig. S12.** Effect sizes from female-biased populations also tend to be from older populations in our dataset. Transparent points represent individual effect sizes scaled by precision (i.e., sample size) and colored according to their effect size. Opaque points and bars represent means +/- 95% CIs.

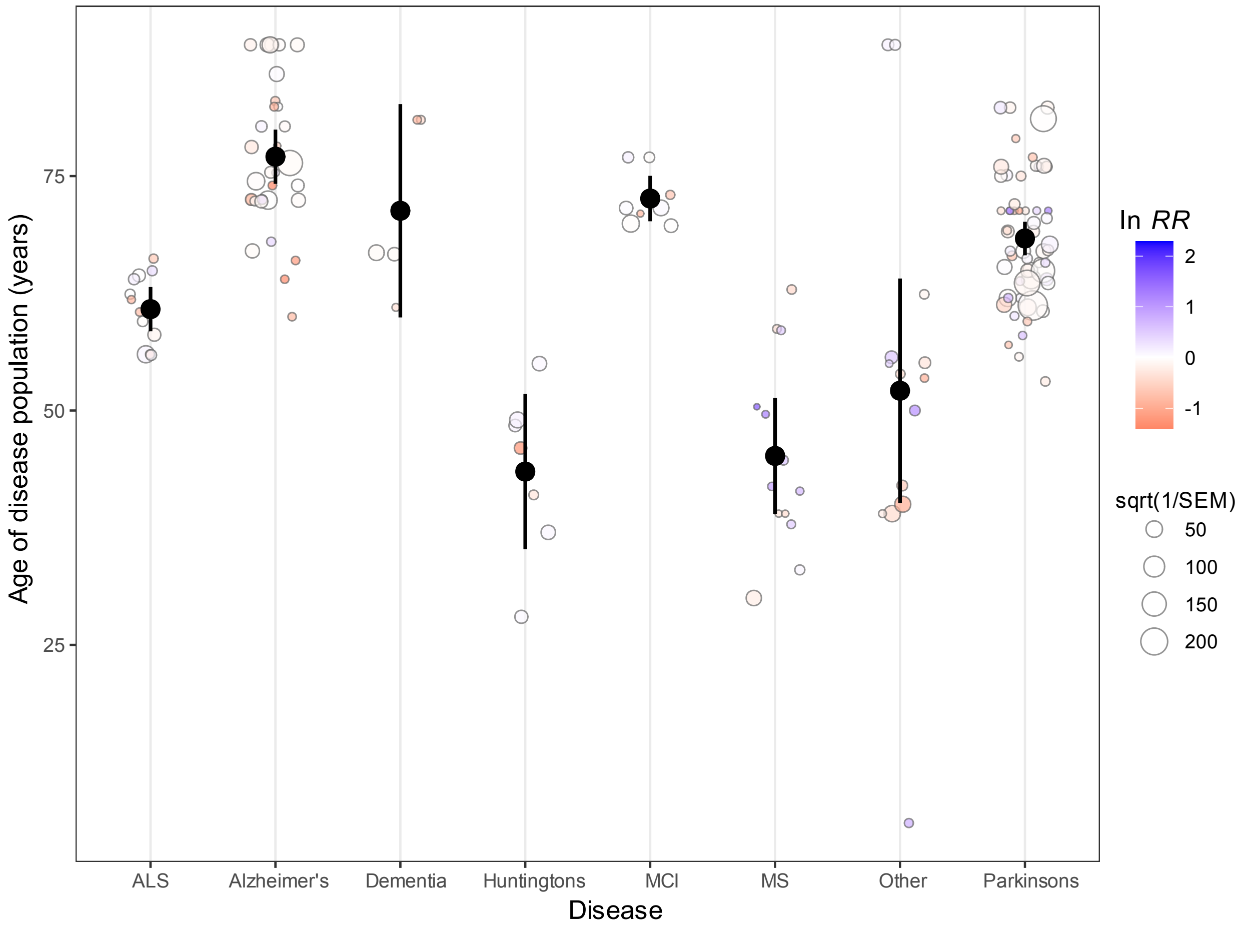


**Fig. S13.** Effect sizes from different diseases have different average ages in our dataset. Transparent points represent individual effect sizes scaled by precision (i.e., sample size) and colored according to their effect size. Opaque points and bars represent means +/- 95% CIs.

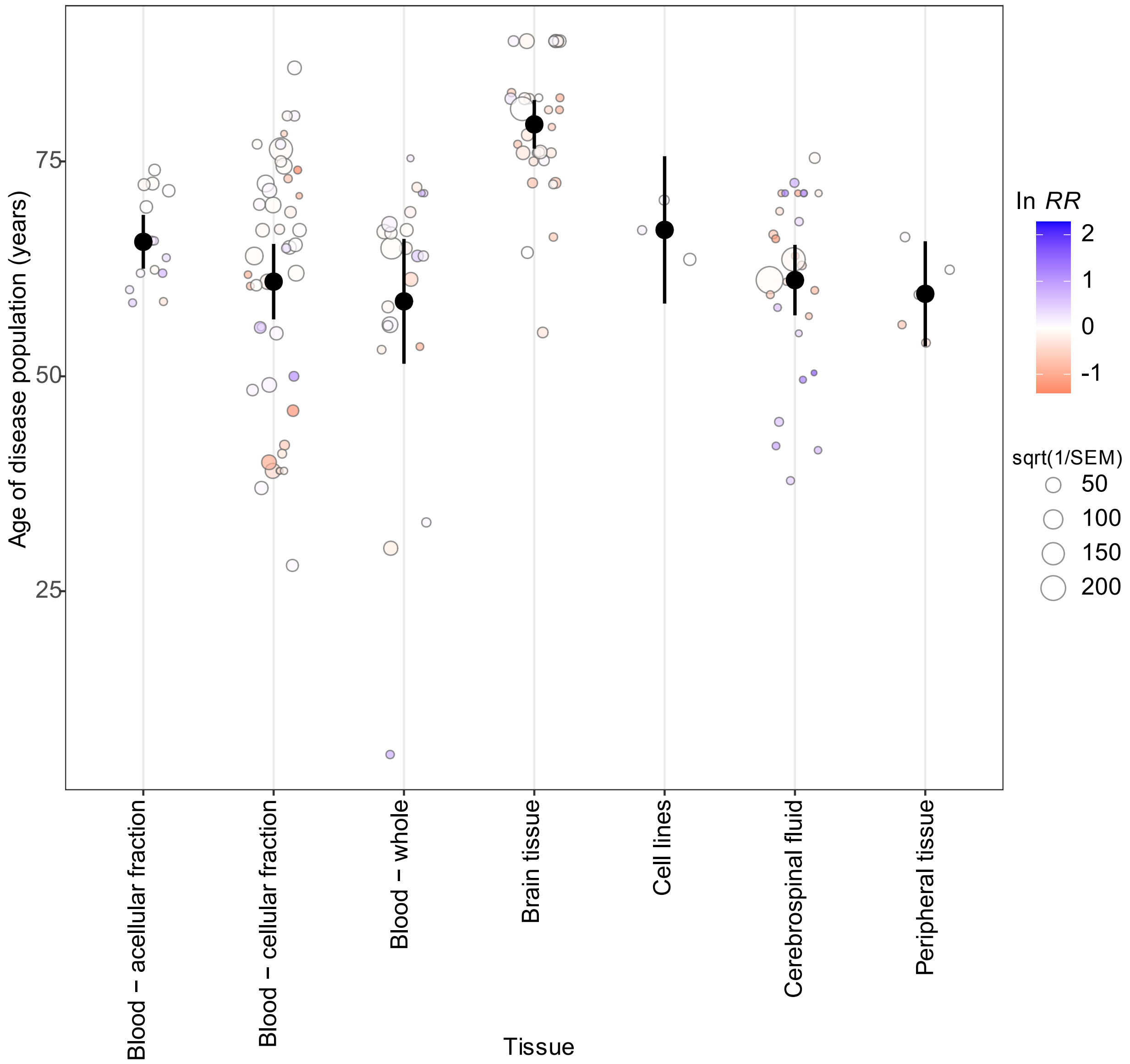


**Fig. S14.** Effect sizes from different tissues have different average ages in our dataset. Transparent points represent individual effect sizes scaled by precision (i.e., sample size) and colored according to their effect size. Opaque points and bars represent means +/- 95% CIs.

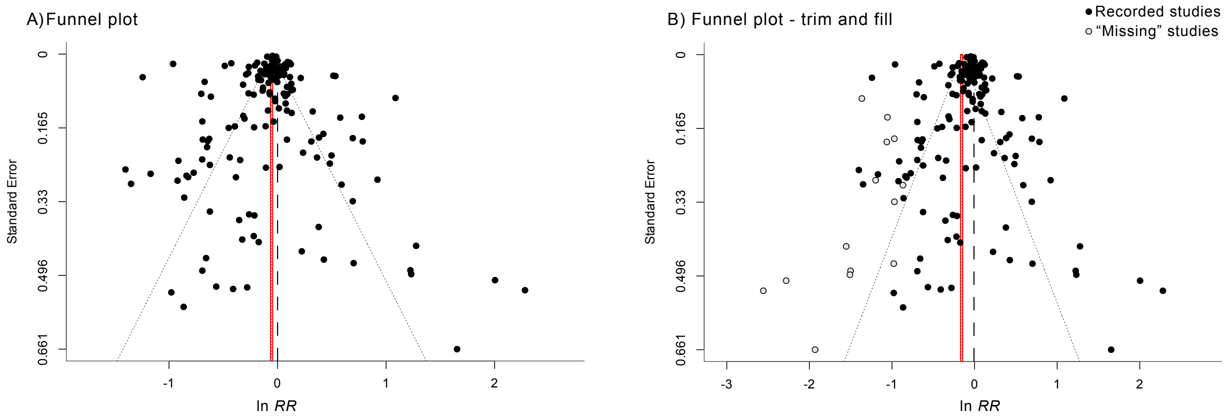


**Fig. S15.** Funnel plots show little evidence of publication bias. A) Traditional funnel plot showing effect sizes plotted against standard error (a measure of precision and sampling power). B) A trim-and-fill funnel plot with "missing" effect sizes (in lighter color) imputed based on funnel plot asymmetry. Red lines indicate grand mean effect sizes (calculated with missing effect sizes in B), dashed line shows no effect (same mtDNA-CN in disease and control population). Symmetrical funnel plots and a relatively low number of missing studies suggest little evidence of publication bias.

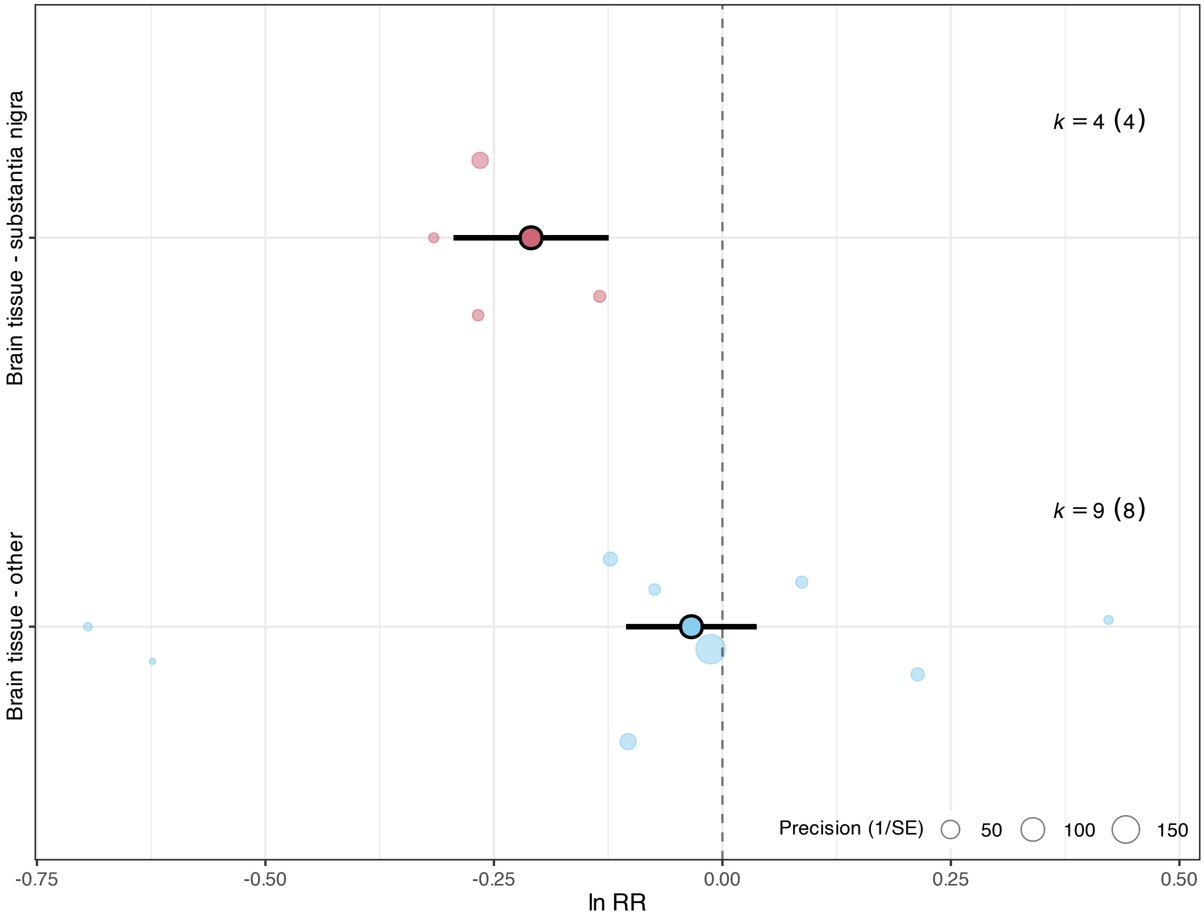


**Fig. S16.** mtDNA-CN is lower with Parkinson's disease in substantia nigra neurons, but is unchanged in other brain regions. Effect sizes are displayed on the x-axis as in Fig. 1, along with *k* values. Transparent points represent individual effect sizes and are scaled to their precision (i.e., sample sizes), while opaque point represent mean effect sizes per category and are shown with 95% CIs.
